# Identification of the Global miR-130a Targetome Reveals a Novel Role for TBL1XR1 in Hematopoietic Stem Cell Self-Renewal and t(8;21) AML

**DOI:** 10.1101/2021.07.30.454489

**Authors:** Gabriela Krivdova, Veronique Voisin, Erwin M Schoof, Sajid A Marhon, Alex Murison, Jessica L McLeod, Martino Gabra, Andy GX Zeng, Eric L Van Nostrand, Stefan Aigner, Alexander A Shishkin, Brian A Yee, Karin G Hermans, Aaron G Trotman-Grant, Nathan Mbong, James A Kennedy, Olga I Gan, Elvin Wagenblast, Daniel D De Carvalho, Leonardo Salmena, Mark D Minden, Gary D Bader, Gene W Yeo, John E Dick, Eric R Lechman

## Abstract

Gene expression profiling and proteome analysis of normal and malignant hematopoietic stem cells (HSC) point to shared core stemness properties. However, discordance between mRNA and protein signatures underscores an important role for post-transcriptional regulation by miRNAs in governing this critical nexus. Here, we identified miR-130a as a regulator of HSC self-renewal and differentiation. Enforced expression of miR-130a impaired B lymphoid differentiation and expanded long-term HSC. Integration of protein mass spectrometry and chimeric AGO2 eCLIP-seq identified TBL1XR1 as a primary miR-130a target, whose loss of function phenocopied miR-130a overexpression. Moreover, we found that miR-130a is highly expressed in t(8;21) AML where it is critical for maintaining the oncogenic molecular program mediated by AML1-ETO. Our study establishes that identification of the comprehensive miRNA targetome within primary cells enables discovery of novel genes and molecular networks underpinning stemness properties of normal and leukemic cells.

**HIGHLIGHTS:**

- miR-130a is a regulator of HSC self-renewal and lineage commitment
- TBL1XR1 is a principal target of miR-130a
- TBL1XR1 loss of function in HSPC phenocopies enforced expression of miR-130a
- Elevated miR-130a levels maintain the AML1-ETO repressive program in t(8;21) AML

## INTRODUCTION

The human blood system is a hierarchically organized continuum with the hematopoietic stem cell (HSC) residing at the apex (Laurenti and Göttgens, 2018; Rieger and Schroeder, 2012; Velten et al., 2017). Hematopoietic homeostasis is tightly regulated by controlling the balance between self-renewal, quiescence and differentiation of HSCs (Laurenti and Göttgens, 2018; Seita and Weissman, 2010). Acquisition of mutations and chromosomal abnormalities in HSC and progenitor cells (HSPC) results in clonal dominance and transformation to acute myeloid leukemia (AML) with leukemia stem cells (LSC) maintaining the malignant hierarchy and driving relapse (Kreso and Dick 2014; Shlush et al., 2014; Thomas and Majeti, 2017; Wang and Dick, 2005). Although transcriptional and epigenetic signatures underpinning different hematopoietic cell states in normal hematopoiesis and leukemia are well described (Buenrostro et al., 2018; Cabezas-Wallscheid et al., 2014; van Galen et al., 2019; Karamitros et al., 2018; Ranzoni et al., 2021; Velten et al., 2017), little is known of the role that post-transcriptional regulation plays in governing the interrelationship between gene expression and proteome signatures of these cell-type specific programs. MicroRNAs (miRNAs) represent a large class of non-coding RNAs that mediate repression of multiple target mRNAs by fine-tuning their expression at the post-transcriptional level. Previous studies indicate that miRNAs are differentially expressed throughout the hematopoietic hierarchy. Moreover, distinct miRNA signatures are associated with specific cytogenetic and molecular subtypes of AML (Garzon et al., 2008; Jongen-Lavrencic et al., 2008; Lechman et al., 2016; Marcucci et al., 2010). Although there is extensive evidence of some miRNAs acting as regulators of stemness and lineage commitment, the mechanism of action and the comprehensive targetome that they repress in HSC are poorly defined. Moreover, how normal miRNA regulatory function is subverted in leukemogenesis and cooperates with specific genetic aberrations present in diverse AML subtypes remains largely unexplored.

In this study, we undertook a focused *in vivo* miRNA enforced expression screen in human HSPC and identified miR-130a as a regulator of stemness and lineage commitment. Our mechanistic studies of the miR-130a targetome revealed repression of gene regulatory networks centered on Transducin Beta-Like 1X-Related Protein 1 (TBL1XR1). TBL1XR1 is a core component of the nuclear corepressor (NCoR) and silencing mediator of retinoic acid and thyroid hormone (SMRT) complexes. Both corepressors form multiprotein complexes consisting of HDAC3, G protein pathway suppressor 2 (GPS2), TBL1XR1 and its closely-related homolog TBL1X (Guenther et al., 2001; Li et al., 2000; Wen et al., 2000; Yoon et al., 2003; Zhang et al., 2002). NCoR and SMRT complexes normally function in repressing transcription governed by unliganded nuclear receptors (NRs) (Chen and Evans, 1995; Hörlein et al., 1995; Perissi et al., 2010; Watson et al., 2012). Binding of steroid hormones such as estrogen and progesterone, or other lipid-soluble factors such as retinoic acid (RA) and thyroid hormone (TR), to their respective receptors initiates dismissal of the corepressor complex and subsequent transcriptional activation (Glass and Rosenfeld, 2000; Perissi et al., 2008; Sever and Glass, 2013). TBL1XR1 and TBL1 act as exchange factors to mediate the dismissal of corepressors and subsequent transcriptional activation of target genes (Perissi et al., 2004, 2008). As NCoR and SMRT complexes mediate and integrate interactions of diverse ligands, receptors and other transcription factors involved in a wide variety of biological processes, including cell proliferation, specification of cell fate and metabolism, it is unsurprising that deregulated function of the repressor complexes has been observed in many types of solid tumors and leukemias (Liang et al., 2019; Wong et al., 2014). However, the function of these complexes across the individual cell types comprising normal and leukemic hematopoietic hierarchies, especially in the context of HSC and LSC, is unknown.

Functional alterations of the NCoR complex have been identified in lymphomas and leukemias with different karyotypic abnormalities. In acute promyelocytic leukemia (APL), which arise from chromosomal translocations generating PML-RARα (promyelocytic leukemia-retinoic acid receptor a) or PLZF-RARα (promyelocytic leukemia zinc finger-retinoic acid receptor α) fusion proteins, increased affinity between the fusion protein and NCoR causes reduced sensitivity to RA-mediated transcriptional activation of target genes involved in myeloid differentiation, thereby promoting leukemogenesis (Atsumi et al., 2006; Grignani et al., 1998; Puccetti and Ruthardt, 2004; Rousselot et al., 1994). Another common translocation in which the function of NCoR is dysregulated is t(8;21) AML. This AML subtype belongs to the core binding factor (CBF) leukemias. RUNX1 (CBF or AML1) is a heterodimeric transcription factor composed of one α (RUNX1, RUNX2, or RUNX3) and one β (CBFβ) subunit (Speck and Gilliland, 2002). RUNX1 functions as a transcriptional activator required for the initial development of definitive hematopoiesis and adult myeloid differentiation (de Bruijn and Speck, 2004; Chen et al., 2009; Guo et al., 2012; Ichikawa et al., 2004; Okuda et al., 1996; Tober et al., 2013). The t(8;21) chromosomal translocation fuses the N-terminal DNA binding domain of the *RUNX1* gene on chromosome 21, to nearly the entire *ETO (RUNX1T1)* gene on chromosome 8 (Erickson et al., 1992, 1994; Miyoshi et al., 1991, 1993). The resulting AML1-ETO onco-fusion protein acts in a dominant negative fashion to inhibit the function of wild-type RUNX1 (Mandoli et al., 2016; Ptasinska et al., 2014; Regha et al., 2015; Stengel et al., 2021). The ETO moiety of AML1-ETO mediates the recruitment of NCoR complex to repress transcription, thereby preventing terminal myeloid differentiation and promoting leukemogenesis (Gelmetti et al., 1998; Link et al., 2010; Lutterbach et al., 1998; Wang et al., 1998). The translocation t(8;21) is a common karyotypic abnormality in AML, accounting for up to 10% of all AML cases (Peterson and Zhang, 2004; Solh et al., 2014). Although this AML subtype is generally associated with a favourable clinical outcome and the rate of complete remission in CBF AML is high (∼87%), relapse-free survival at 10 years following standard therapy is only 30-50%, indicating that recurrence and relapse-related death remain a challenge in the treatment of this heterogenous disease (Appelbaum et al., 2006; Marcucci et al., 2005). Thus, a better understanding of the molecular mechanisms driving t(8;21) AML initiation, progression and maintenance is critical for the development of targeted therapies and improved clinical outcomes.

The ability of miRNAs to repress multiple targets resulting in downregulation of various signalling pathways underscores the importance of uncovering their complete targetome in order to understand the mechanistic function of miRNAs in normal hematopoiesis and clinical significance of dysregulated expression in AML. To our knowledge, this is the first study to utilize chimeric AGO2 eCLIP in human HSPC and an AML cell line to uncover endogenous miRNA-target interactions. By integrating miRNA-target chimeras with proteomics and RNA-seq data, we establish the mechanistic function of miR-130a in human HSPC and t(8;21) AML. Our findings reveal that repression of TBL1XR1 by miR-130a in normal HSPC impedes differentiation and expands the functional LT-HSC population. Moreover, our study demonstrates that elevated miR-130a levels in t(8;21) AML facilitate leukemogenesis by reinforcing the repressive AML1-ETO molecular program and preventing differentiation of leukemic cells.

## RESULTS

### *In vivo* microRNA Screen Identifies miR-130a as a Regulator of HSC Function

To investigate the functional role of candidate miRNAs previously found to be differentially expressed in normal and malignant human HSPC (Lechman et al., 2016), we performed a competitive repopulation screen to determine the impact of the overexpression (OE) of individual miRNAs on long-term repopulation potential. Lineage-negative (Lin^-^) CD34^+^ CD38^-^ cord blood (CB) cells were transduced with individual lentiviral constructs encoding selected miRNAs and mOrange^+^(mO^+^) reporter gene and transplanted into the right femur of immune-deficient NOD/Lt-scid/IL2RD percentag^null^ (NSG) mice (Figure 1A). Competitive repopulation was assessed by the fold change in the transduced human cells (mO^+^/hCD45^+^) at 24 weeks following transplantation over mO^+^ input levels. Enforced expression of miR-125b, miR-155 and miR-130a significantly increased long-term hematopoietic reconstitution, consistent with the previously reported role of miR-125b and miR-155 in the self-renewal of murine HSC (Itkin et al., 2012; Ooi et al., 2010). In contrast, miR-10a and miR-196b OE significantly reduced the repopulation capacity of HSPC (Figure 1B, Figure S1A). Enforced expression of several miRNAs altered lineage distribution, including increased erythroid lineage output with miR-155 and miR-130a OE (Figure S1B-C). Based on the observed increase in long-term competitive repopulation and altered lineage output of human HSPC with miR-130a OE, we prioritized this miRNA for functional validation studies.

**Figure 1.**
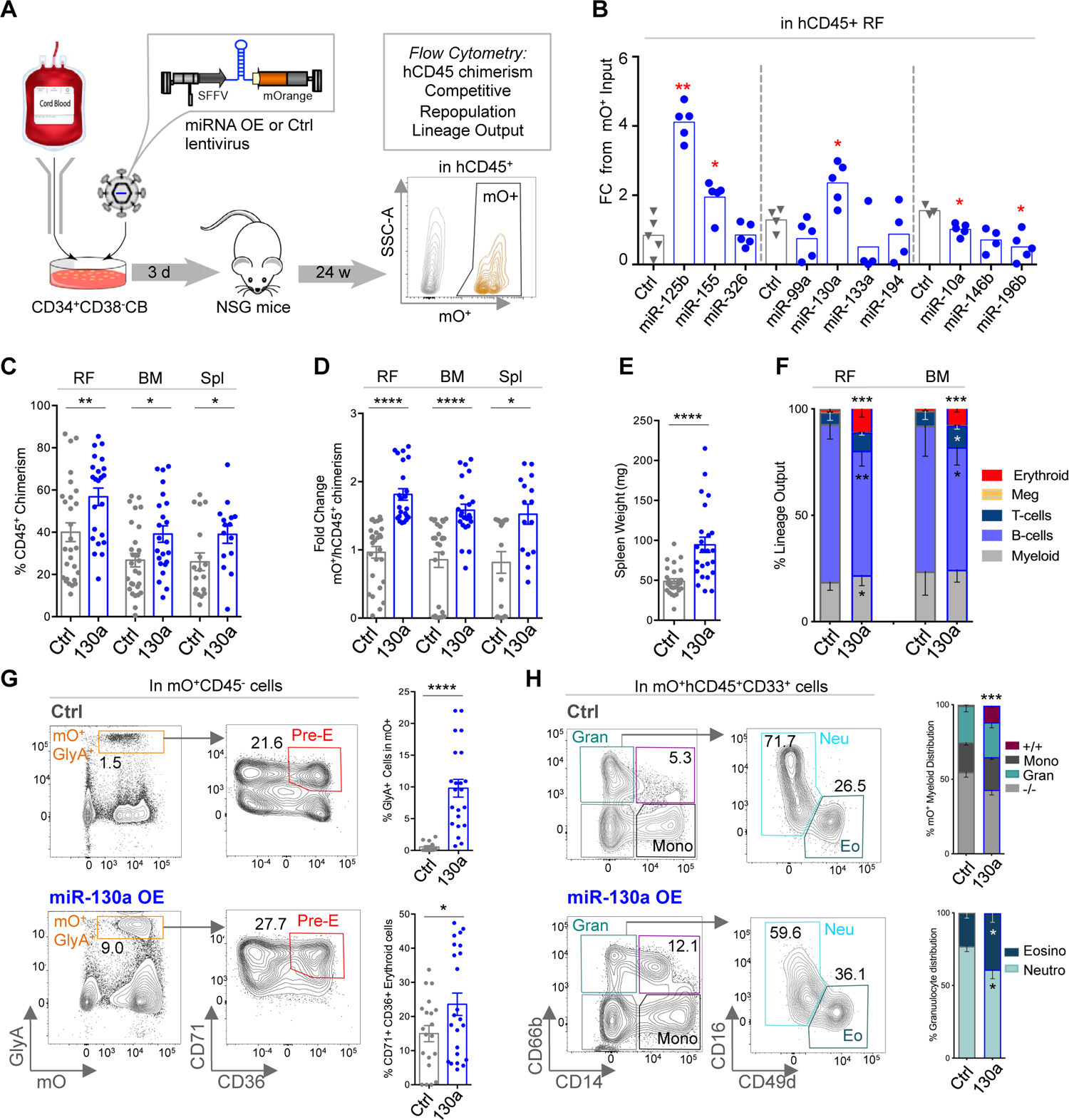
Enforced Expression of miR-130a Enhances Long-Term Repopulation Capacity and Alters Lineage Output of HSC. (A) Schematic of the *in* vivo miRNA overexpression (OE) repopulation assay. (B) Fold change of mO^+^CD45^+^ cells in right femur (RF) at 24 weeks post-transplantation compared to input levels following enforced expression of individual miRNAs in HSPC. (C) Human CD45^+^ chimerism in right femur (RF), bone marrow (BM) and Spleen (Spl) at 24 weeks post-transplantation with HSPC transduced with miR-130a OE or control lentiviruses (n=3 biological experiments, 7-10 mice/experimental group). (D) Fold change of mO^+^CD45^+^ cells at 24 weeks post-transplantation compared to input levels. (E) Spleen weight of xenotransplanted mice. (F) Lineage distribution of mO^+^ xenografts. Graphs reflect mean values from 3 independent biological experiments, 7-10 mice/experimental group. (G) Representative flow cytometry plots of GlyA^+^ erythroid cells and CD36^+^CD71^+^ progenitors (left) and graphs presenting proportion of these populations in mO^+^GlyA^+^ cells (right). (H) Representative flow cytometry plots of CD14^+^ monocytes, CD66b^+^ granulocytes, CD49d^+^ eosinophils and CD16^+^ neutrophils (left) and graphs presenting proportion of these populations in mO^+^hCD45^+^CD33^+^ myeloid cells (right). (B-H) Mann-Whitney test, all error bars indicate ± SEM, *p<0.05, **p<0.01, ***p<0.001, ****p<0.0001.

To investigate the impact of miR-130a OE on human HSC function, we transduced Lin-CD34^+^ CD38^-^ CB cells with miR-130a OE or control lentivectors and assessed engraftment by flow cytometry 24 weeks post-xenotransplantation. Increased expression of miR-130a in human mO^+^CD45^+^ cells from xenografts was confirmed by RT-qPCR (Figure S1D). Enforced expression of miR-130a resulted in greater human chimerism in the injected right femur (RF), non-injected bone marrow (BM) and spleen compared to controls (Figure 1C). Notably, miR-130a OE conferred a competitive advantage over non-transduced cells and resulted in enlarged spleens of the recipient mice (Figure 1D, 1E). Examination of the lineage distribution of mO^+^ cells revealed significant increases in erythroid, myeloid and T cell output at the expense of B-cell differentiation (Figure 1F, S1E-I). Of note, miR-130a OE xenografts exhibited accumulation of immature CD71^+^CD36^+^ erythroid progenitors, suggestive of a block in erythroid differentiation (Figure 1G). Enforced expression of miR-130a also altered the myeloid differentiation program as evidenced by the presence of an aberrant CD33^+^ cell population co-expressing granulocytic CD66b and monocytic CD14 markers (Figure 1H) along with significantly higher proportions of eosinophils and reduced levels of neutrophils within miR-130a OE granulocytes (Figure 1H). To further assess the functional significance of miR-130a in human HSPC, we used a knockdown strategy (KD) utilizing a miRNA sponge lentivector with 8 tandem copies of imperfectly complementary miR-130a target sequence designed to specifically inhibit miR-130a function and GFP as a reporter gene. miR-130a KD caused substantial changes in lineage differentiation that were opposite to our observed OE phenotype, including significantly increased B-lymphoid lineage output at the expense of myeloid and erythroid differentiation (Figure S1J-P). We also observed decreased proportions of CD36^+^ cells in GlyA^+^ erythroid cells and expansion of CD14^+^ monocytes at the expense of granulocytic differentiation (Figure S1O, S1P). Collectively, our data show that enforced expression of miR-130a increases engraftment potential of human HSPC, impairs B-lymphoid lineage output and alters erythroid and myeloid differentiation programs.

### Enforced Expression of miR-130a Expands Functional HSC

Given the observed effect of miR-130a on long-term repopulation and lineage output, we next sought to identify the HSPC populations in which miR-130a exerts its function. Combined BM and RF cells harvested from engrafted mice were depleted of murine and human lineage-committed cells and the remaining HSPC were analyzed by flow cytometry. Enforced expression of miR-130a resulted in an approximately 2-fold increase in the proportion of CD34^+^CD38^-^ cells and an increased proportion of immunophenotypic LT-HSC (CD34^+^CD38^-^CD45RA^-^CD90^+^) (Figure 2A, 2B). The proportions of more committed progenitors within the CD34^+^CD38^+^ compartment were not altered (Figure 2C). The absolute number of CD34^+^CD38^-^ and CD34^+^CD38^+^ cells was increased 11-fold and 6-fold, respectively, in miR-130a OE xenografts compared to control (Figure S2A). Secondary transplantation at limiting dilution demonstrated an approximately 10-fold higher frequency of LT-HSC (1 in 1.5×10^5 cells) in recipients transplanted with miR-130a OE HSPC compared to controls (Figure 2D, S2B). In contrast, miR-130a KD caused notable reduction of the proportion of GFP^+^ cells in the Lin-compartment and a decrease in the number of CD34^+^CD38^-^ and CD34^+^CD38^+^ cells (Figure S2C). Together, these results implicate a role for miR-130a in enhancing HSC self-renewal.

**Figure 2.**
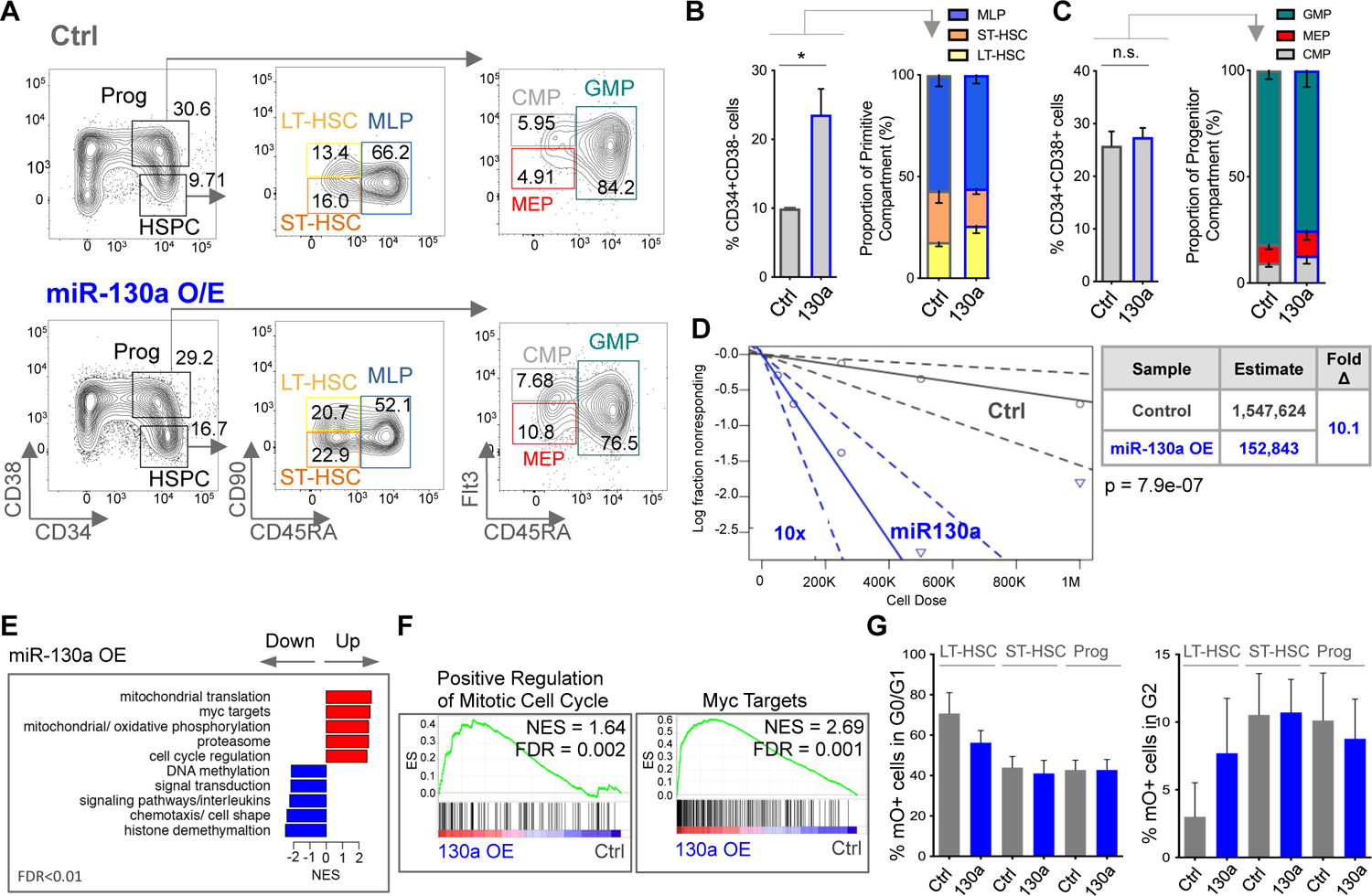
Enforced Expression of miR-130a Expands HSC by Forcing Them Into the Cell Cycle. (A) Representative flow plots of HSPC populations from control and miR-130a OE xenografts at 12 weeks. (B) Proportion of CD34^+^CD38^-^ compartment and frequency of LT-HSC, ST-HSC and MLP cell populations (n=4, each replicate contains pooled RF and BM from 2-4 individual mice). (C) Proportion of CD34^+^CD38^+^ compartment and frequency of CMP, MEP and GMP cell populations. (D) Secondary transplantation of CD45^+^mO^+^ from 12 weeks NSG mice at limiting dilution doses. Frequency of HSC was evaluated from the engraftment in secondary mice at 12 weeks from two independent biological experiments (>0.05% CD45^+^mO^+^ cells in BM, n=50). (E) Normalized enrichment score (NES) of the gene sets significantly different between miR-130a OE and control-transduced CD34^+^ HSCP (n=3). (F) GSEA plots of HSC cell cycle primed gene sets and MYC targets in transcriptome profile following miR-130a OE in CD34^+^ HSPC. (G) Cell cycle analysis of LT-HSC, ST-HSC and CD34^+^CD38^+^ progenitor cells transduced with miR-130a or control lentiviruses (n=3) (B-C) Unpaired t-test, all error bars indicate ± SEM, *p<0.05.

To identify the molecular pathways that contribute to the enhanced self-renewal and competitive repopulation advantage observed in miR-130a OE HSPC, we performed RNA-seq on mO^+^CD34^+^ HSPC 3 days after transduction with control or miR-130a OE lentiviral constructs (Table S1). Gene set enrichment analysis (GSEA) revealed enrichment of mitochondrial translation, MYC signalling, proteasome function and cell cycle regulation pathways in genes upregulated in miR-130a OE cells (Figure 2E, S2D). In contrast, histone demethylation, interleukin signalling and DNA methylation pathways were enriched in genes downregulated upon miR-130a OE (Figure 2E, S2D). As many of the upregulated gene sets following miR-130a OE were related to HSC programs, we performed GSEA using quiescent and activated HSC signatures (Forsberg et al., 2010) and observed enrichment of mitotic cell cycle activation and proliferation gene sets and depletion of quiescent genes with miR-130a OE (Figure 2F, S2E). We also observed significant enrichment of MYC targets following miR-130a OE (Figure 2F), consistent with HSC activation and quiescence exit (García-Prat et al., 2021; Laurenti et al., 2008). To determine whether miR-130a alters proliferation and cell cycle kinetics of HSC, we performed EdU incorporation assays utilizing fluorescently-labelled thymidine analog to measure nascent DNA synthesis in sorted LT-HSC and ST-HSC 3 days after transduction with control or miR-130a lentiviruses. Enforced expression of miR-130a in LT-HSC but not ST-HSC resulted in a decrease in the proportion of mO^+^ cells in G0/G1 phase and an increase in G2 phase, corroborating a role for miR-130a in cell cycle activation (Figure 2G, S2F). Thus, enforced expression of miR-130a induces HSC activation and cell cycle progression while expanding the functional HSC pool.

### Repression of TBL1XR1 by miR-130a Contributes to Downregulation of Chromatin Remodelling and Lipid Metabolism Pathways

To characterize potential miR-130a targets, we performed low cell input, quantitative, label-free mass spectrometry analysis (Schoof et al., 2016) in control or miR-130a OE CD34^+^ CB cells (Table S2). GSEA revealed 145 upregulated and 128 downregulated pathways following miR-130a OE at a p value <0.05 (Figure S1A). As miRNAs generally act as negative regulators of gene expression, we focused our analysis of miR-130a targets on downregulated proteins. To identify which downregulated proteins are potential targets of miR-130a, we integrated experimentally-validated Tarbase targets into the subsequent analysis. Tarbase miR-130a targets were strongly enriched among proteins downregulated upon miR-130a OE at a p value <0.05 (Figure S3B, S3C). Enrichment mapping revealed that chromatin modification, interferon signaling and lipid metabolism were significantly enriched in both downregulated proteins and miR-130a Tarbase targets (Figure 3A, S3D). Examination of overlap between predicted (mirDIP) and experimentally-validated (Tarbase) miR-130a targets identified 9 genes that were also downregulated in our proteomics dataset, including TBL1XR1, CBFβ and JARID2 (Figure 3B, S3E). Interestingly, JARID2 and CBFβ are components of the well-characterized Polycomb Repressive Complex 2 (PRC2) and RUNX1 complexes that regulate chromatin organization and gene expression programs essential for definitive hematopoiesis and HSC function and are frequently mutated in leukemia (Cai et al., 2015; Ichikawa et al., 2004; Majewski et al., 2008; Radulović et al., 2013; Xie et al., 2014). Repression of TBL1XR1, CBFβ and JARID2 by miR-130a OE was validated by size-based, capillary western blot in human CD34^+^ CB cells ( gure 3C, Figure S3E). Enrichment mapping of the 9 repressed miR-130a targets with the downregulated gene sets revealed association of TBL1XR1 with multiple gene networks including chromatin organization/modification, HDAC function and lipid metabolism (Figure 3D).

**Figure 3.**
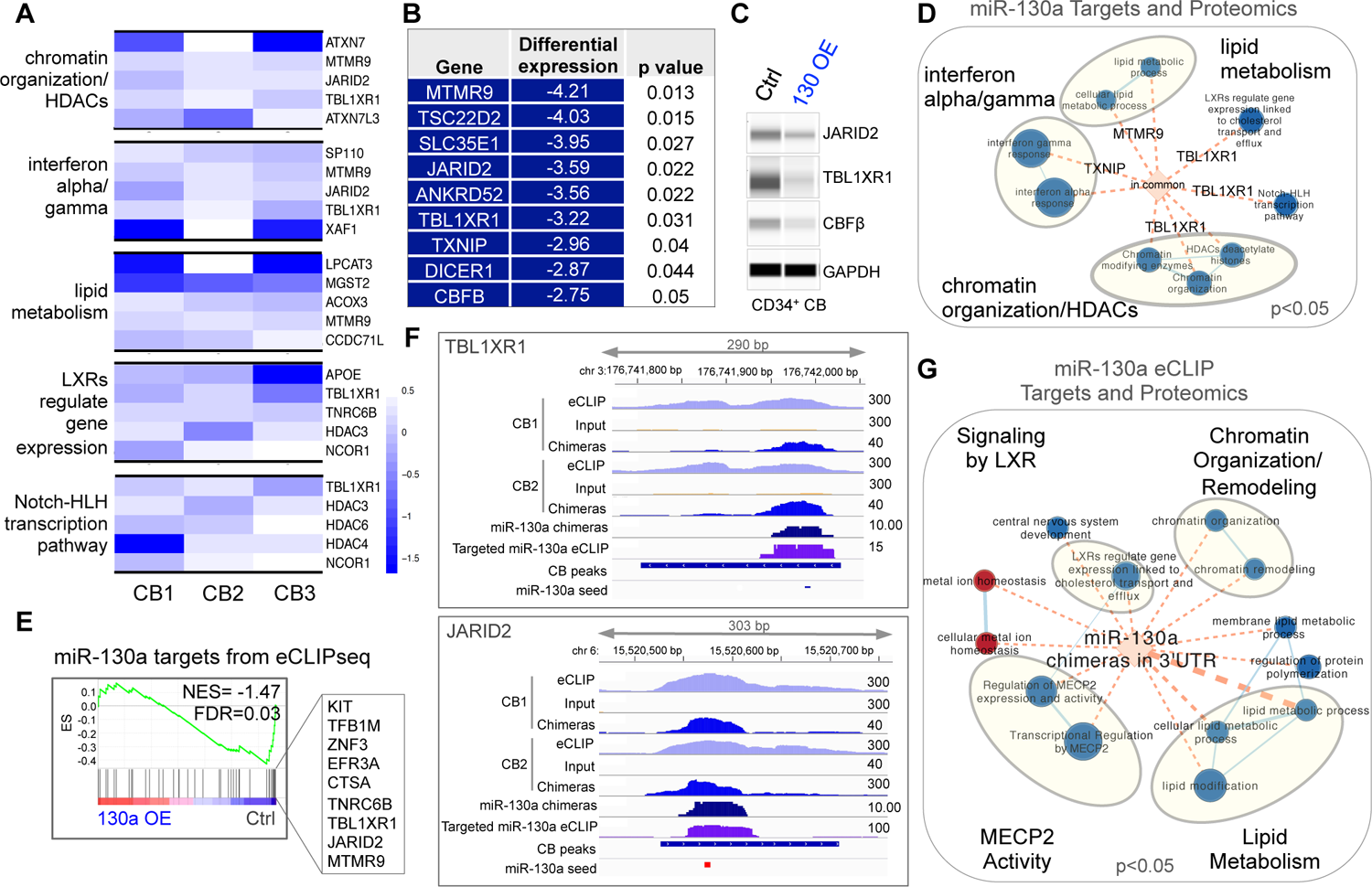
Mass Spectrometry and Chimeric AGO2 eCLIP Reveal miR-130a Targetome in Human HSPC. (A) Heat map of downregulated proteins and experimentally supported TarBase targets showing downregulated pathways (left side) and selected proteins (right side) following miR-130a OE in CD34^+^ CB cells; Wilcoxon one-sided test, p<0.05. (B) List of miR-130a downregulated proteins that are predicted TarBase and mirDIP targets, limma t-test. (C) Western blot showing repression of miR-130a targets in mO^+^CD34^+^CB. (D) Enrichment map showing gene sets containing miR-130a targets listed in B; FDR<0.05, node size is proportional to NES; Wilcoxon one-sided test, p<0.05. (E) GSEA plots of miR-130a targets from chimeric AGO2 eCLIP in proteome profile following miR-130a OE in CD34^+^ HSPC and a list of leading edge genes. (F) Genome browser tracks presenting all reads from AGO eCLIP libraries, size-matched input control, chimeric AGO2 eCLIP reads, chimeric reads containing only miR-130a sequence and reads from targeted miR-130a eCLIP. eCLIP peaks represent reproducible regions of enrichment from both CB replicates normalized over input controls. (G) Enrichment map of miR-130a targets from chimeric AGO2 eCLIP in downregulated protein sets following miR-130a OE; FDR<0.05, node size is proportional to NES, Wilcoxon one-sided test; p <0.05.

Currently, the only biochemical method to identify direct miRNA-target interaction sites *in vivo* is by UV crosslinking and purification of Argonaute (AGO)-bound RNA complexes (Chi et al., 2009; Darnell, 2010; Grosswendt et al., 2014; Helwak et al., 2013; Moore et al., 2015). To capture endogenous miR-130a targets in human HSPC, we performed a modified AGO2 enhanced cross-linking and immunoprecipitation (eCLIP)-seq technique (Van Nostrand et al., 2016) in CD34^+^ CB, and analyzed chimeric reads containing miRNAs ligated to their targets (Table S3). Additionally, due to the requirement of the eCLIP method for large input cell numbers, we developed a targeted eCLIP-seq method which incorporates PCR amplification using miRNA-specific primers for more sensitive detection of miRNA-target interactions. Analysis of miR-130a-target chimeras revealed 57% of miR-130a targets detected by eCLIP were present in Tarbase (Figure S3G). As miRNA target sites located in the 3’UTR of genes were previously shown to be associated with the greatest target repression (Broughton et al., 2016; Grimson et al., 2007; Grosswendt et al., 2014; Helwak et al., 2013), we selected miR-130a chimeras with 3’UTR target sites for further analysis. We focused on chimeras that were present in both targeted and transcriptome-wide eCLIP-seq experiments (n= 67) (Table S4). GSEA revealed significant enrichment of miR-130a-target chimeras in the list of proteins downregulated by miR-130a OE in comparison with control at a p value <0.05 (Figure 3E). Notably, we identified MTMR9, TBL1XR1, JARID among the leading edge genes enriched in the downregulated proteins (Figure 3E, S3G). Genome browser tracks containing total and chimeric miR-130a eCLIP-seq reads showed peaks corresponding to binding of miR-130a to the 3’UTR of TBL1XR1, JARID2 and CBFβ, providing evidence for direct binding of miR-130a to these transcripts (Figure 3F, S3H). Intersection of the miR-130a target chimeras with the downregulated proteins showed enrichment of lipid metabolism and chromatin remodelling/organization and signaling by liver X receptors (LXR), in line with our mass spectrometry data (Figure 3G). Intriguingly, we noted that methyl CpG binding protein (MeCP) activity was also significantly downregulated following miR-130a OE.

MECP2 directly interacts with TBL1XR1, allowing its recruitment to heterochromatin and association with the NCoR and SMRT complexes (Kruusvee et al., 2017). Moreover, we observed downregulation of LXR activity, which belongs to the family of nuclear receptors (NR), with concomitant decrease in protein levels of NCoR and HDACs, suggesting that enforced expression of miR-130a impairs signaling via the NR/NCoR pathway. Collectively, integration of mass spectrometry and chimeric AGO2 eCLIP-seq techniques identified TBL1XR1 as a direct miR-130a target that contributes to downregulation of genes involved in chromatin organization/modification, interferon signaling, lipid metabolism and NR signaling pathways.

### Repression of TBLXR1 by miR-130a Impedes Differentiation and Expands LT-HSC

Since we observed downregulation of genes involved in the NR signaling pathway including TBL1XR1, which is required for transcriptional activation of several NR including retinoic acid receptor (RAR), we interrogated expression levels of RA-target genes following miR-130a OE. GSEA revealed depletion of genes activated by RA in miR-130a OE CD34^+^ CB cells (Figure S4A). As the function of TBL1XR1 in hematopoiesis and HSC is largely unknown, we investigated the impact of TBL1XR1 deficiency in human HSPC by performing KD studies in human CB-derived CD34^+^ HSPC. We tested 4 TBL1XR1 shRNAs for knockdown efficiency and selected 2 shRNAs for *in vivo* studies (Figure S4B). Western blotting confirmed efficient KD of TBL1XR1 in transduced cells from xenografts marked by BFP reporter gene (Figure 4A). TBL1XR1 KD significantly decreased human chimerism measured by the percentage of human CD45^+^ cells and the proportion of BFP^+^ cells in the injected RF and non-injected BM of mice at 24 weeks post-transplantation (Figure 4B, 4C). Lineage analysis of TBL1XR1 KD xenografts revealed a significant decrease in mature CD19^+^ B cells and an increase in CD33^+^ myeloid cells and CD45^+^ human cells that did not express committed lineage markers (Lin-) (Figure 4D, S4C-G). Within the myeloid lineage, we observed a significant decrease in mature granulocytes and an increase in monocytes in TBL1XR1 KD xenografts (Figure S4H). Analysis of the HSPC compartment revealed an increase in CD34^+^CD38^-^ cells and significant expansion of immunophenotypic LT-HSC and ST-HSC populations with a concomitant decrease in the MLP population (Figure 4E, 4F). We also observed a significant expansion of CD7^-^CD10^-^ progenitors out of the CD34^+^CD38^+^ population and an increase in the proportion of myelo-erythroid progenitors upon TBL1XR1 KD (Figure S4I). Serial transplantation at limiting dilution of BFP^+^CD45^+^ cells harvested at 24 weeks from primary recipients into secondary NSG-GF recipients, which allows for a more sensitive readout of HSC frequency, demonstrated a 4.7 and 2.6-fold increase in HSC frequency with TBL1XR1 shRNA3 and shRNA4, respectively (Figure 4G, S4J). In line with our findings from primary xenografts, TBL1XR1 KD significantly impaired B cell differentiation in secondary recipients, resulting in 80-90% myeloid engraftment (Figure S4K). Thus, TBL1XR1 KD in human HSPC functionally phenocopies enforced expression of miR-130a in expanding LT-HSC and impairing B-lymphoid differentiation.

**Figure 4.**
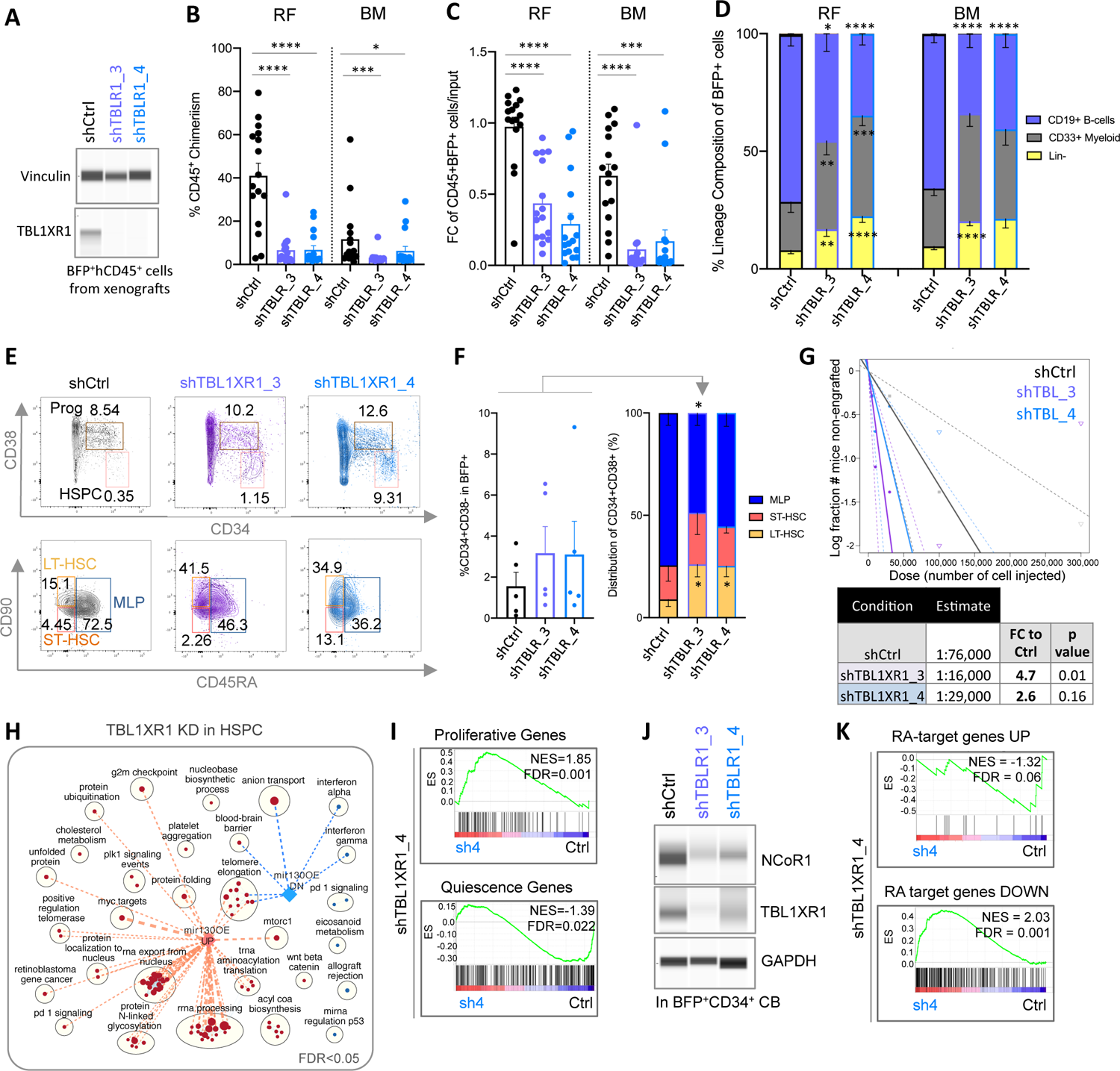
Repression of TBL1XR1 Impairs B-cell Differentiation and Expands LT-HSC. (A) Western blot showing TBL1XR1 protein levels in CD45^+^BFP^+^ cells from 12 week xenografts from control and TBL1XR1 KD mice. (B) Human CD45^+^ chimerism in right femur (RF) and bone marrow (BM) at 24 weeks post-transplantation with HSPC transduced with two TBL1XR1 shRNAs (sh3 and sh4) or control shRNA lentiviruses (n=2 biological experiments, 6-8 mice per experimental group). (C) Fold change of BFP^+^CD45^+^ cells in RF and BM at 24 weeks post-transplantation compared to input levels. (D) Lineage distribution of BFP^+^ xenografts. (E) Flow cytometry plot of HSPC populations from control and TBL1XR1 KD xenografts. Flow plots are representative of triplicate samples overlaid together. (F) Proportion of BFP^+^CD34^+^CD38^-^ cells in RF and frequency of LT-HSC, ST-HSC and MLP cell populations in 24 week xenografts from two independent biological experiments (n=5, each replicate contains pooled RF from 2-4 individual mice), unpaired t-test. (G) Secondary transplantation of CD45^+^BFP^+^ from 24 week NSG mice at limiting dilution doses. Frequency of HSC was evaluated from the engraftment in NSG-GF secondary mice at 8 weeks (>0.05% CD45^+^BFP^+^cell in BM, n=39). (H) Enrichment map of upregulated (red nodes) and downregulated (blue nodes) pathways following TBL1XR1 KD in CD34^+^CD38^-^ CB cells intersected with miR-130a-target chimeras from chimeric AGO2 eCLIP, FDR<0.05, hypergeometric t-test, p<0.05. (I) GSEA plots of proliferative and quiescence genes in transcriptome profile following TBL1XR1 KD. (J) Western blot showing protein levels of NCoR and TBL1XR1 following TBL1XR1 KD in CD34^+^ CB cells. (K) GSEA plots of genes upregulated and downregulated by retinoic acid in transcriptome profile following TBL1XR1 KD. (B-D) Unpaired Mann-Whitney u-test, all error bars indicate ± SEM, *p<0.05, **p<0.01, ***p<0.001.

To investigate the molecular mechanism of HSC expansion associated with TBL1XR1 KD, we performed RNA-seq of BFP^+^ progeny of transduced CD34^+^CD38^-^ HSPC cultured *in vitro* (Table S1). Interestingly, we observed a significant decrease in the expression of TBL1XR1 and a concomitant increase in its homolog TBL1X at FDR 0.05 (Figure S4L). Despite the increased expression level of the TBL1X homolog, we observed significant gene expression changes upon TBL1XR1 KD, arguing against redundancy between the two factors in human HSPC (Table S1). Enrichment mapping revealed upregulation of genes in rRNA processing pathways and MYC targets in TBL1XR1 KD HSPC, concordant with the observed enrichment of these gene sets following enforced expression of miR-130a (Figure 4H). Similar to the gene expression changes observed with miR-130a OE, GSEA showed enrichment of cell cycle activation and proliferation genes and depletion of genes associated with cellular quiescence with TBL1XR1 loss of function (Figure 4I, S4M). As TBL1XR1 is a core component of the NCoR complex involved in transcriptional repression and activation of nuclear receptor signaling pathways, we performed western blot analysis of TBLXR1-deficient CD34^+^ CB cells and confirmed decreased level of NCoR (Figure 4J). Moreover, analysis of RA-target genes, which are known to be regulated by TBL1XR1 and NCoR (Li and Wang, 2008; Perissi et al., 2004), revealed strong enrichment of genes repressed by RA and depletion of genes upregulated by RA in TBL1XR1 KD cells, implicating that a consequence of TBL1XR1 deficiency is abrogation of repression with concomitant activation of the RA-target genes (Figure 4K). Overall, these findings demonstrate that repression of the miR-130a target TBL1XR1 largely phenocopies the functional effects of enforced expression of miR-130a with concomitant changes in gene expression including upregulation of translation and cell cycle activation pathways.

### High miR-130a Expression is Required for Maintenance of t(8;21) AML

Our data identified miR-130a as a regulator of stemness and lineage specification in normal human HSPC. As blocks in myeloid differentiation and aberrant self-renewal programs are characteristic of leukemia and several identified miR-130a targets are involved in oncogenic pathways in AML, we investigated whether expression of miR-130a is deregulated in some AML subtypes. Using the TCGA dataset, we identified four AML clusters based on miR-130a expression (Figure S5A). AML subtypes expressing high levels of miR-130a had higher LSC17 scores, a stemness signature associated with adverse risk in AML (Ng et al., 2016) (Figure S5B). As an independent validation of higher miR130a levels in AML cells reflecting increased stemness properties, we calculated the maximum enrichment of chromatin accessibility over background at the miR-130a locus in our ATAC-seq data from functionally-defined LSC^+^ and LSC^-^ AML fractions and observed increased chromatin accessibility surrounding the miR-130a locus in LSC^+^ fractions compared to LSC^-^ fractions at p value <0.05 (Figure S5C). Next, we mapped miR-130a expression onto TCGA, BEAT-AML and Leucegene AMLs classified on the basis of leukemic hierarchy composition into granulocyte-monocyte progenitors (GMP), intermediate, mature and primitive groups using our new CIBERSORTx-based deconvolution approach involving single-cell RNAseq signatures (Zeng et al. submitted). We observed high expression of miR-130a in primitive and GMP-like AMLs compared to mature AMLs (Figure 5A). Similarly, GSVA using single cell RNA-seq data of normal hematopoietic populations revealed association of miR-130a expression with cells exhibiting high HSC-progenitor and GMP-like signatures and low myeloid-like signatures (Figure S5D). Of note, the GMP-like AML cluster is highly enriched for CBF-rearranged AMLs including t(8;21) and inv(16); these subtypes showed significantly higher levels of miR-130a compared to other AML subtypes in the TCGA dataset (Figure 5A, 5B). Moreover, the majority of t(8;21) and inv(16) AML samples segregated into the clusters with high miR-130a expression (Figure S5E). To determine if miR-130a is of clinical importance, we examined miR-130a levels in CBF AML patient samples obtained from Princess Margaret Hospital Leukemia Tissue Bank (Table S5) and observed that high miR-130a levels were associated with shorter overall and disease-free survival (Figure 5C, Figure S5F). RT-qPCR analysis of miR-130a expression levels in CD34^+^ blasts from an independent cohort of CBF AML patients from PM (TableS5) revealed higher miR-130a expression compared to CD34^-^ leukemic cells and approximately 4-fold higher miR-130a levels compared to peripheral blood mononuclear cells from healthy controls (Figure 5D). In addition, profiling of miR-130a expression across different AML cell lines showed significantly higher expression in the Kasumi-1 cell line, which is derived from a t(8;21) AML and is characterized by enrichment for the GMP-like hierarchy signature (Figure S5G, S5H). Collectively, these data show that miR-130a is highly expressed in CBF AML, including t(8;21) AML, and that higher miR-130a expression identifies a subset of CBF patients with inferior outcome.

**Figure 5.**
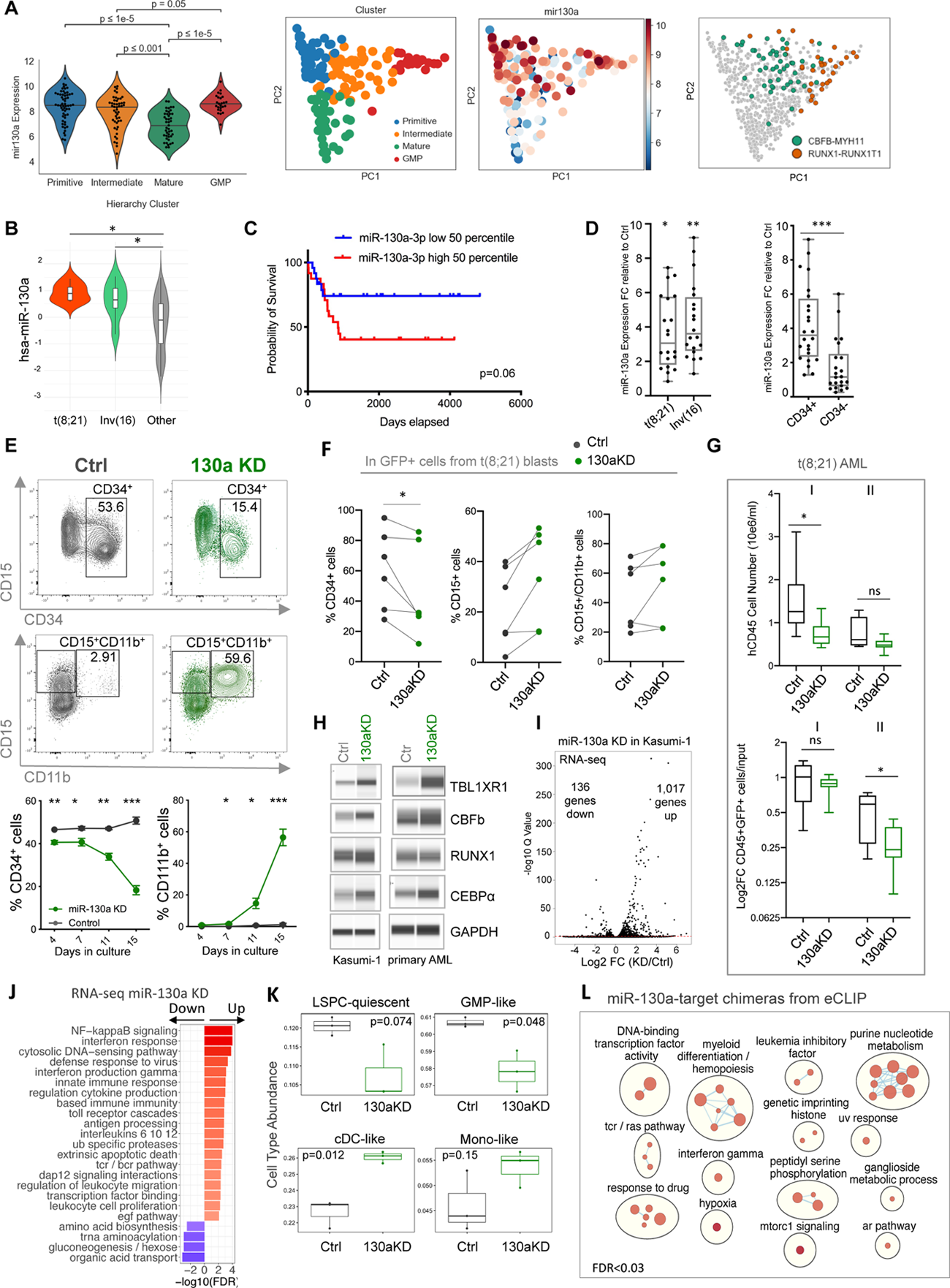
Loss of Function of miR-130a Induces Differentiation of t(8;21) AML. (A) Clustering of AML subtypes into 4 groups based on scRNA-seq data; expression of miR-130a and position of CBF AML in these clusters are shown. (B) Expression of miR-130a in t(8;21) and inv(16) AML compared to other subtypes from the TCGA dataset. (C) Kaplain-Meir curve showing correlation between miR-130a expression (50% split) and CBF AML patient survival. (D) qRT-PCR showing expression of miR-130a in CD34^+^ blasts from t(8;21) AML compared to healthy controls (n=20) and CD34^-^ cells from the same patient samples (n=24). (E) Flow cytometry plots and graphs (below) representing immunophenotype of Kasumi-1 cells transduced with control or miR-130a KD lentiviruses (n=3). (F) Graphs representing quantitative changes in CD34, CD15 and CD11b surface markers expression on t(8;21) AML blasts 8 days post transductionwith control and miR-130a KD lentiviruses. (G) Graphs representing changes in CD45^+^ leukemic engraftment and log2 fold change of GFP^+^ control or miR-130a KD cells 10 weeks post-transplantation compared to input conditions in two primary t(8;21) samples in NSG-GF mice. Unpaired Mann-Whitney u-test, all error bars indicate ± SEM, *p<0.05. (H) Western blot showing levels of TBL1XR1, CBFβ, RUNX1 and CEBP following miR-130a KD in Kasumi-1 cells. (I) Volcano plot showing global gene expression changes following miR-130a KD in Kasumi-1 cells. (J) Bar graph representing upregulated and downregulated gene sets following miR-130a KD in Kasumi-1 cells, FDR<0.01. (K) Deconvolution of gene expression changes of control and miR-130a KD Kasumi-1 cells based on the AML signatures from the scRNA-seq data. (L) Enrichment map of upregulated gene sets following miR-130a KD in Kasumi-1 overlaid with miR-130a targets from chimeric AGO2 eCLIP, FDR<0.03, Mann-Whitney, p<0.05. (A-K) Unpaired t-test unless indicated otherwise., all error bars indicate ± SEM, *p<0.05, **p<0.01, ***p<0.001.

To examine the functional significance of high miR-130a expression in t(8;21) AML, we performed loss of function studies in Kasumi-1 cells and primary AML patient samples. In Kasumi-1 cells, miR-130a KD induced a significant reduction in the CD34^+^ cell population and an increase in differentiated CD11b^+^CD15^+^ cells and annexinV^+^ apoptotic cells over a period of 2 weeks in culture (Figure 5E, S5I). Similarly, miR-130a KD in CD34^+^ blasts from six t(8;21) AML patient samples followed by culture on MS5 stroma cell line expressing human colony-stimulating factor 1 (hCSF1) resulted in a decreased proportion of CD34^+^ cells and an increased percentage of differentiated CD11b^+^CD15^+^ cells compared to control (Figure 5F, S5J-K). Transplantation into NSG-GF mice of two t(8;21) patient samples transduced with miR-130a KD lentivector resulted in lower engraftment and a smaller proportion of GFP^+^ cells compared to controls (Figure 5G). miR-130a KD in Kasumi-1 cells and primary t(8;21) AML cells led to increased levels of TB1XR1 and CBFβ detected by western blot, consistent with de-repression of these targets (Figure 5H). We also detected increased levels of the myeloid differentiation transcription factor CEBPα, consistent with the differentiation immunophenotype observed with miR-130a KD (Figure 5H). Together, these results demonstrate that elevated miR-130a expression is required for maintenance of t(8;21) leukemia and its loss of function results in differentiation and apoptosis of leukemic cells.

To examine the effect of miR-130a on global gene expression in Kasumi-1 cells, we performed RNA-seq of Kasumi-1 cells following miR-130a KD (Table S1). We observed global activation of gene expression, including 1,017 transcripts upregulated and 136 transcripts downregulated following miR-130a KD at FDR <0.05 (Figure 5I). Genes in interferon/immune response, ubiquitin processing/degradation and apoptosis/cell proliferation pathways were strongly enriched following miR-130a KD (Figure 5J), in line with our functional data showing a differentiation phenotype and apoptosis. Deconvolution of Kasumi-1 cells using the CIBERSORTx approach revealed loss of LSPC-quiescent and GMP-like signatures and a gain in cDC-like and Mono-like signatures upon miR-130a KD (Figure 5K), consistent with the observed functional requirement of miR-130a in the differentiation block and maintenance of t(8;21) AML. To elucidate which of the upregulated transcripts are direct miR-130a targets, we performed chimeric AGO2 eCLIP-seq in Kasumi-1 cells and identified 100 and 890 miR-130a-target chimeras from the global and targeted approaches, respectively (Table S4). miR-130a chimeric reads containing the 3’UTR of TBL1XR1 and CBFβ were captured by the more sensitive, targeted eCLIP-seq approach. By focusing on targets with binding located in the 3’UTR and shared chimeras between the targeted and global approaches, we narrowed the list of miR-130a direct targets to 40 genes including JARID2, TXNIP, TNRC6B and KMT2C. Examination of the overlap between the miR-130a chimeras and the enrichment map revealed association of these miR-130a targets with upregulated pathways, confirming global de-repression of targets following miR-130a KD (Figure S5L). Myeloid differentiation and interferon signaling pathways were enriched in genes upregulated in miR-130 chimeras (Figure 5L). Collectively, our results indicate that miR-130a KD in t(8;21) AML causes de-repression of targets centered on myeloid differentiation and interferon responses, leading to upregulation of inflammatory pathways and apoptosis.

### miR-130a Maintains the Repressive AML1-ETO Gene Network by Governing its Composition and Binding

Apart from the interactions with NCoR and HDAC, AML1-ETO associates with several hematopoietic transcription (co)factors including CBFb, RUNX1, ERG, E proteins HEB and E2A, LMO2 and LYL1, which contributes to leukemogenesis (Mandoli et al., 2016; Sun et al., 2013). Given that miR-130a is highly expressed in t(8;21) AML and our identification of TBL1XR1 and CBFβ as direct miR-130a targets in normal HSPC, we performed immunoprecipitation (IP) of AML1-ETO in Flag tag-knock-in AML1-ETO (Flag-AE) Kasumi-1 cells to interrogate changes in the composition of the AML1-ETO complex following miR-130a KD. The specificity of the Flag antibody and Flag-AML1-ETO IP was confirmed with a wild–type (WT) Kasumi-1 cell line and a control IP sample with no addition of Flag peptide for elution (Figure S6A, S6B). Western blot analysis revealed increased levels of CBFβ and TBL1XR1 and decreased levels of LMO1 and HEB in the input lysate in miR-130a KD cells. Surprisingly, we observed decreased association of AML1-ETO with CBFβ in the miR-130a KD cells compared to control in the IP sample (Figure 6A). Moreover, western blot analysis revealed loss of association of AML1-ETO with NCoR and the transcription factors HEB and LMO2 (Figure 6A), suggesting that high miR-130a levels are required to maintain the composition of the AML-ETO complex.

**Figure 6.**
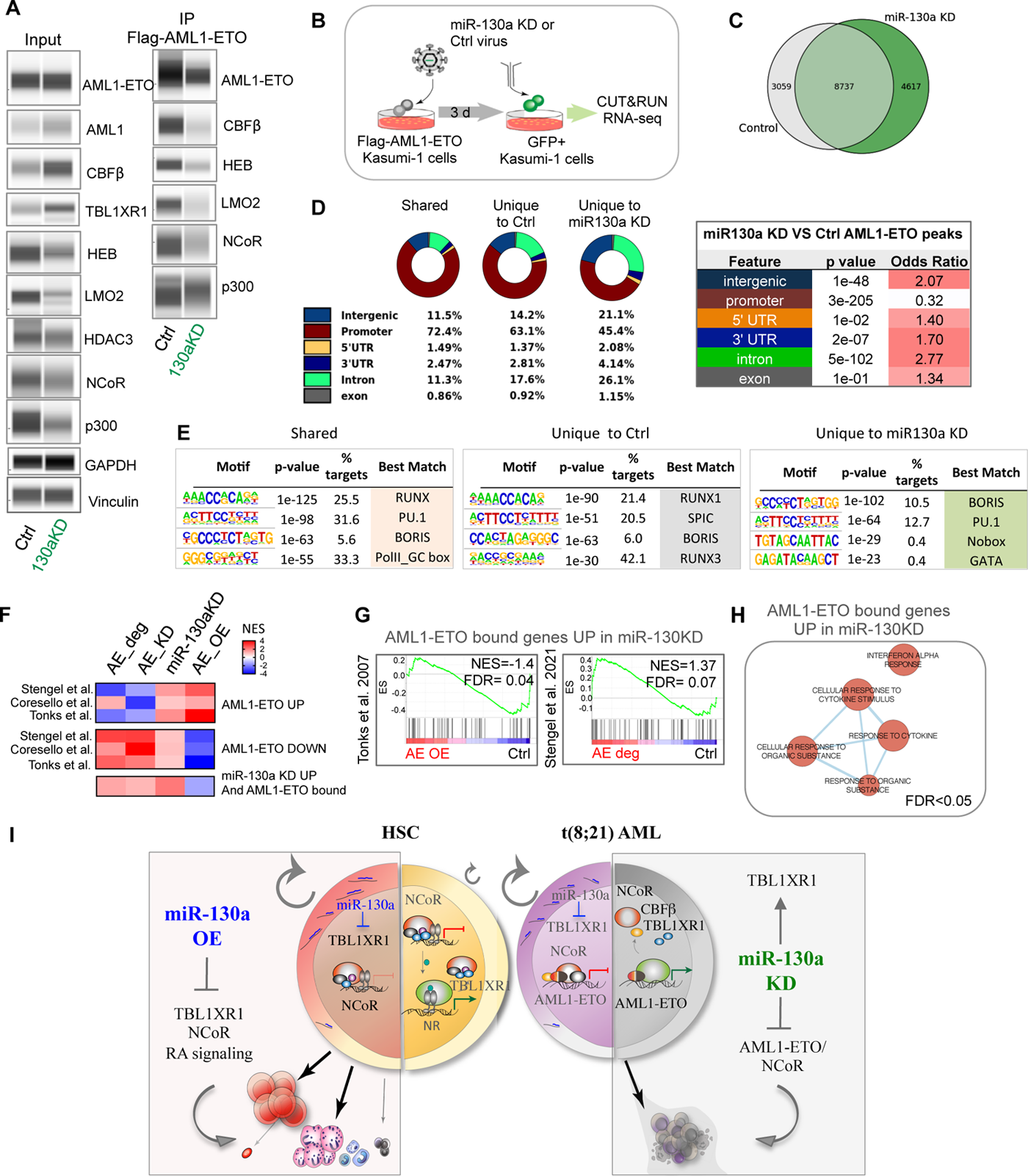
miR-130a Controls the Composition and Binding of the AML1-ETO Complex. (A) Western blots showing input lysates and IP of Flag-AML1-ETO in Kasumi-1 cells transduced with control and miR-130a KD lentiviruses. (B) Schematic depiction of the experimental design for RNA-seq and CUT&RUN assay following miR-130a KD in Flag-tagged AML1-ETO Kasumi-1 cells. (C) Venn diagram showing shared and unique AML1-ETO peaks in control and miR-130a KD Flag-AML1-ETO Kasumi-1 cells (q<0.001). (D) Genomic distribution of AML1-ETO peaks in control and miR-130a KD Kasumi-1 cells (left). Fisher T-test was used to compare the distribution of peaks between control and miR-130a KD (right). (E) HOMER transcription factor binding site motif enrichment analysis of shared and unique peaks from control and miR-130a KD Kasumi-1 cells. (F) Heat map of gene expression signatures from AML1-ETO degradation, knock-down and overexpression datasets compared to transcriptome profile following miR-130a KD in Kasumi-1 cells. (G) GSEA plots showing enrichment of genes bound and upregulated by AML1-ETO in miR-130a KD Kasumi-1 cells in gene expression changes following AML1-ETO overexpression (left) or degradation (right). (H) Enrichment map showing upregulated, promoter-bound AML1-ETO genes unique to miR-130a KD; genes FDR<0.05, pathways FDR<0.05. (K) Mechanistic model of miR-130a function in normal HSC and t(8;21) leukemia.

To investigate whether changes in the composition of the AML1-ETO complex following miR-130a KD alter its binding occupancy, we performed CUT&RUN (cleavage under targets and release using nuclease) and RNA-seq (Figure 6B). CUT&RUN assays were performed with anti-Flag and anti-ETO antibodies in Flag-AML1-ETO Kasumi-1 cells, as the Kasumi-1 cells do not express ETO. In control cells, 12,260 of the 13,816 peaks (89%) detected with ETO antibody were also detected with anti-Flag antibody, demonstrating a substantial overlap between the binding sites (Figure S6C). In contrast, the overlap between peaks detected by anti-Flag and anti-ETO antibodies was less pronounced (62%) in miR-130a KD cells (Figure S6C). Importantly, the majority (∼85%) of our annotated AML1-ETO binding sites overlapped with published CUT&RUN datasets of HA-tagged AML1-ETO in Kasumi-1 cells (Stengel et al., 2021) (Figure S6D). Analysis of shared peaks captured with both anti-Flag and anti-ETO antibodies, herein referred to as AML1-ETO peaks, revealed 4,617 unique peaks in miR-130a KD, 3,059 unique peaks in control and 8,737 peaks shared between control and miR-130a KD Kasumi-1 cells (Figure 6C). Interestingly, AML1-ETO binding regions unique to miR-130a KD cells displayed significant loss of binding sites in promoter regions (45%) compared to peaks unique to control (63.1%) and shared peaks (72.4%) as well as a gain in intronic and intergenic regions at p<0.001 (Figure 6D). Motif enrichment analysis of the peaks mapping to promoter regions of AML1-ETO-bound genes revealed the RUNX motif to be the most highly enriched motif in both the shared peaks and as well as peaks unique to control (Figure 6E, S6E), supporting the specificity of the CUT&RUN approach. In contrast, RUNX motifs were not enriched in peaks unique to miR-130a KD cells (Figure 6E, S6E), implicating a shift in AML1-ETO binding with a resultant loss of specificity. K-means clustering based on intensity of peaks unique to control identified 2 clusters, both with high enrichment of the RUNX motif (43% and 23%) (Figure S6F). In contrast, AML1-ETO peaks unique to miR-130a KD segregated to 3 clusters, none of which showed a strong enrichment of the RUNX motif (Figure S6F). Gene ontology (GO) analysis of promoter-bound AML1-ETO genes unique to control showed enrichment in cell cycle and DNA damage pathways. In comparison, lipid metabolism, MAPK and translation pathways were enriched in AML1-ETO promoter-bound regions unique to miR-130a KD cells (Figure S6G). Overall, these findings show that loss of function of miR-130a in t(8;21) AML results in reorganization of the AML1-ETO repressor complex, including loss of association with CBFβ and NCoR, with resultant altered binding of AML1-ETO across the genomic landscape.

Previous studies have reported that expression of AML1-ETO results mainly in transcriptional repression of direct gene targets (Ptasinska et al., 2014, 2019; Stengel et al., 2021). Degradation of AML1-ETO in Kasumi cells results in de-repression of direct gene targets constituting a core AML1-ETO regulatory network, whereas AML1-ETO overexpression in HSPC causes repression of these gene targets (Stengel et al., 2021; Tonks et al., 2007). To assess the impact of miR-130a KD on this known AML1-ETO gene network, we integrated RNA-seq analysis with AML1-ETO binding occupancy from the CUT&RUN analysis and interrogated several RNA-seq and ChIP-seq datasets including AML1-ETO OE in HSPC and AML1-ETO KD and degradation in Kasumi-1 cells (Corsello et al., 2009; Stengel et al., 2021; Tonks et al., 2007). AML1-ETO target genes identified from our CUT&RUN analysis as being promoter-bound and also upregulated following miR-130a KD were enriched in the AML1-ETO degradation and KD gene sets (Figure 6F, 6G). These same genes were depleted in AML1-ETO OE samples, implying these AML1-ETO targets are de-repressed following miR-130a loss of function (Figure 6F, 6G). This set of genes was strongly enriched in interferon alpha and cytokine response pathways (Figure 6H, S6H) and transcription factor enrichment analysis using iRegulon predicted STAT proteins as regulators of these pathways (Figure S6I-K). Collectively, our findings reveal a unique role of miR-130a in regulating normal hematopoietic stem cell self-renewal and how elevated levels of miR-130a in t(8;21) AML contribute to the leukemogenesis of this AML subtype (Figure 6I).

## DISCUSSION

Our study establishes a fundamental mechanistic role for miR-130a in human HSCs. Using an *in vivo* competitive repopulation screen and gain/loss of function assays, we established that miR-130a is a regulator of HSC self-renewal and lineage commitment, with its enforced expression severely impairing B-lymphoid differentiation and expanding LT-HSC. To better understand how miR-130a governs these properties within HSC, we integrated *in vivo* miRNA enforced expression and loss of function studies with chimeric AGO2 eCLIP-seq to capture endogenous miRNA targetomes in physiologically relevant primary cells. TBL1XR1 was identified as a principal miR-130a target, whose repression results in downregulation of chromatin organization, lipid metabolism and NR pathways controlled by NCoR. TBL1XR1 loss of function largely phenocopied miR-130a enforced expression, establishing TBL1XR1 as a novel regulator of HSC self-renewal and lineage differentiation and underscoring the role of NR signaling pathways in human HSC. Moreover, our study provides insight into how elevated miR-130 levels maintain the AML1-ETO repressive transcriptional program in t(8;21) AML by governing the composition and binding occupancy of the AML-ETO/NCoR repressor complex. Overall, these data provide evidence of the essential role that miRNA-mediated post-transcriptional regulation serves within both normal and malignant hematopoietic stem cells.

Our findings demonstrate that enforced expression of miR-130a in human HSPC promotes cell cycle entry and activates MYC and translation pathways without compromising HSC integrity, resulting in expansion of the functional LT-HSC population. Although several studies had shown that miR-130a is highly expressed in HSC and myeloid progenitors and that its expression declines during differentiation (Gentner et al., 2010; Georgantas et al., 2007; O’Connell et al., 2010; Petriv et al., 2010), the function of miR-130a in human HSC was previously unknown. Our characterization of the global miR-130a targetome in CD34^+^ HSPC revealed repression of several known hematopoietic factors including CBFβ and JARID2, and, importantly, identified TBL1XR1 as a new regulator of HSC function. Our analysis suggests that TBL1XR1 is associated with the preponderance of pathways that are downregulated in HSPC following miR-130a OE including chromatin organization, lipid metabolism, LXR and MeCP2 function. In support of our results, independent studies in non-hematological tissues show that MeCP2 directly interacts with TBL1XR1 and NCoR/SMRT repressor complexes (Kruusvee et al., 2017) Mutations in MeCP2 or TBL1XR1 are common in neurological and developmental disorders and disrupt MeCP2 interaction with NCoR, suppressing the ability NCoR to repress transcription (Ebert et al., 2013; Kruusvee et al., 2017; Lyst et al., 2013). Moreover, signaling via LXR is tightly regulated by TBL1XR1, whereby depletion of TBL1XR1 reduces ligand dependent LXR activation of key target genes involved in cholesterol homeostasis in liver cells (Jakobsson et al., 2009), underscoring the significance of TBL1XR1 and NCoR in orchestrating NR signalling pathways across different tissues.

Although NR pathways and the NCoR/SMRT complexes are generally well studied in development and tissue homeostasis using bulk tissues and tissue-specific mutant animal models (Feng et al., 2001; Jepsen et al., 2000; Koide et al., 2001; Mottis et al., 2013; Xu et al., 2009), their function within the heterogeneous cell types that make up tissue hierarchies, including the role they may play in regulating stem cell self-renewal and fate specification, remains largely unexplored. Only a limited number of studies have addressed the contribution of NCoR/SMRT complexes and diverse NR pathways in regulating HSC function. Conditional knockout of NCoR1 in murine LSK cells severely impaired B-cell differentiation and increased frequency of LT-HSC, ST-HSC, multipotent progenitor and myelo-erythroid progenitor (Wan et al., 2019). Cell cycle analysis of NCoR-deficient LT-HSC and murine HSPC (LSK) cells revealed significant reduction in G0 cells with concomitant increases in G1 and S/G2/M cell cycle phases, implicating NCoR in regulation of HSC quiescence and proliferation (Wan et al. 2019). Our study corroborates the role of NcoR controling HSC properties and uncovers a novel function of TBL1XR1, a core component of the NCoR/SMRT repressor complexes, in regulating HSC self-renewal and cell fate specification. Our data point to a convergent function of NCoR and TBL1XR1 in promoting lymphoid differentiation and HSC activation and further underscore the under-studied role of NR pathways in regulating stemness properties of HSC. TBL1XR1 is highly homologous with two related genes, TBL1RX and TBL1RY. Previous reports suggest that while thyroid receptor (TR), estrogen receptor (ER), peroxisome proliferator-activated receptor (PPAR) and NF-kB-mediated transcriptional activation requires both TBL1XR1 and TBL1X, RAR and AP-1-mediated activation is exclusively dependent on TBL1XR1 (Li and Wang, 2008; Perissi et al., 2004). Although we detected the homolog TBL1X to be among the top transcripts upregulated following TBL1XR1 KD, our results argue against functional redundancy of the two factors as reported previously in other tissues (Yoon et al., 2005). A similar TBL1X upregulation following TBL1XR1 loss of function was observed in neuronal progenitor cells, which exhibited defective proliferation, implying an exclusive role of TBL1XR1 in a subset of signalling cascades associated with NCoR (Mastrototaro et al., 2021). Our results show that TBL1XR1 KD abrogates signalling by RA, demonstrated by de-repression of targets downregulated by RA and the inability to activate RA target genes in TBL1XR1-deficient cells. As RA signalling has been shown to regulate HSC maintenance/function (Cabezas-Wallscheid et al., 2017; Ghiaur et al., 2013) and TBL1XR1 is required for the transcriptional activation by NRs, we speculate that the inability to activate pathways downstream of NR including RA and impaired function of the NCoR/SMRT complexes contributes to HSC activation and enhanced self-renewal of TBL1XR1-deficient HSC. Investigation of the precise role of TBL1XR1, TBL1X and NCoR-mediated repression and activation of signalling pathways governed by NRs in purified human HSC subsets will be challenging due to limited cell numbers, but clearly warranted. Additionally, since miRNAs act by repressing multiple targets, we expect that downregulation of other miR-130a targets including JARID2, MTMR9 and TXNIP, contribute to the observed phenotype following enforced expression of miR-130a in HSPC.

Our data demonstrate that high miR-130a levels in t(8;21) AML are required for leukemia maintenance. Indeed, previous miRNA profiling studies reported elevated expression of miR-130a in CBF-rearranged leukemias including t(8;21) and inv(16) (Ding et al., 2018; Li et al., 2008). Moreover, miR-130a expression was shown to be elevated at relapse compared to diagnosis and associated with worse event-free survival of t(8;21) AML (Ding et al., 2018). Here, we show that miR-130a loss of function induces differentiation and apoptosis of t(8;21) AML cells accompanied by upregulation of genes enriched in myeloid differentiation and interferon signalling, suggesting that high miR-130a levels are required for t(8;21) AML maintenance. Moreover, our findings from immunoprecipitation of AML1-ETO and CUT&RUN experiments revealed altered composition and chromatin binding of the AML1-ETO complex following miR-130a KD and de-repression of AML-ETO target genes implicating that miR-130a is required for the maintenance of the AML1-ETO oncogenic program. Decreased association of CBFβ and NCoR with AML1-ETO implies that the repressor complex composition is subverted by miR-130a loss of function. We hypothesize that reduced levels of TBL1XR1 mediated by high miR-130a levels contribute to the proper assembly and functioning of the AML1-ETO/NCoR complex. Interestingly, focal deletions and recurrent point mutations of TBL1XR1 commonly occur in pediatric B-ALL and are enriched at relapse compared to diagnosis, implying that reduction in TBL1XR1 activity contributes to leukemogenesis (Mullighan et al., 2008; Parker et al., 2008; Zhang et al., 2011). Focal deletions of TBL1XR1 were reported in 10-15% of B-ALL patients with the translocation t(12;21) characterized by the oncofusion protein ETV6-RUNX which also recruits NCoR to mediate repression. Decreased expression of TBL1XR1 in B-ALL was associated with increased levels of NCoR and resistance to glucocorticoid drugs (Jones et al., 2014). Moreover, recurrent fusions involving TBL1XR1 occur in acute promyelocytic leukemia and involve translocation between TBL1XR1, RARA and RARB genes resulting in diminished transcriptional activity of RAR and inhibition of myeloid maturation (Chen et al., 2014; Osumi et al., 2018, 2019). Collectively, these studies and our findings point to a novel function of TBL1XR1 in normal hematopoiesis and implicate that its loss of function contribute to diverse hematologial maligancies. Our study describing the mechanistic function of miR-130a in t(8;21) provide a novel insight into the complexity of the interactions between the different factors that constitute the AML1-ETO/NCoR repressive complex. The mechanism driving upregulated expression of miR-130a in this AML subtype is unclear and further understanding of how its expression is regulated could enable therapeutic manipulation of miR-130a expression levels to induce differentiation of t(8;21) AML. Moreover, our data reveal that miR-130a is also highly expressed in inv(16) AML characterized by the fusion protein CBFb-SMMHC with high levels being associated with inferior overall survival; however, the functional significance of elevated expression of miR-130a in this AML subtype is unclear and warrants further investigation. Collectively, our characterization of miR-130a function and identification of its comprehensive targetome in normal HSPC and t(8;21) AML has shed light on the transcriptional control of the pathways mediated by TBL1XR1 and NCoR in normal hematopoiesis and AML. Further investigation of the complex interactions between AML1-ETO, NCoR and TBL1XR1 will be beneficial for the design of future differentiation therapies targeting the repressor complex with the goal of developing better outcomes for patients with t(8;21) leukemia.

## Acknowledgments

We thank the obstetrics units at Trillium, William Osler and Credit Valley hospitals for providing the cord blood samples; the Animal Resource Centre (UHN) for support with the mouse work; SickKids UHN Flow and Princess Margaret Flow Facility for FACS; the Center for Applied Genomics and Princess Margaret Genomics Centre for next generation sequencing and the laboratories of Steven Chan and Faiyaz Notta for sharing equipment. We thank Scott W. Hiebert for supplying the Flag-tagged AML1-ETO Kasumi-1 cell line; Steven Henikoff for Protein A-Micrococcal nuclease and spike-in yeast DNA and M.D.M. for the human CSF1 MS5 stroma cell line. We thank Jean C.Y. Wang for detailed editing the manuscript. We thank members of the Dick lab for assistance and critical feedback; Scott W. Hiebert, M.D.M, Courtney Jones and Anastasia Tikhonova for comments on the manuscript.

## Funding

G.K. is supported by Canadian Institutes of Health Research (CIHR) Doctoral Award, Frederick Banting and Charles Best Canada Graduate Scholarship (CGS-D). This work to J.E.D. was supported by funds from the: Princess Margaret Cancer Centre Foundation, Ontario Institute for Cancer Research through funding provided by the Government of Ontario, Canadian Institutes for Health Research (RN380110 - 409786), International Development Research Centre Ottawa Canada, Canadian Cancer Society (grant #703212 (end date 2019), #706662 (end date 2025)), Terry Fox New Frontiers Program Project Grant (Project# 1106), University of Toronto’s Medicine by Design initiative with funding from the Canada First Research Excellence Fund, a Canada Research Chair, Princess Margaret Cancer Centre, The Princess Margaret Cancer Foundation, and Ontario Ministry of Health. This work was supported by NIH grant HG009889 to G.W.Y.

## Author Contributions

G.K. conceived the study, performed *in vitro* and *in vivo* experiments, analyzed data and wrote the manuscript. V.V. performed downstream analysis of RNA-seq, MS and eCLIP-seq data, including enrichment mapping, GSEA and analysis of TCGA AML. E.M.S. performed and analyzed mass spectrometry data. S.A.M. performed CUT&RUN analysis. A.M. performed RNA-seq analysis. J.L.M., E.R.L., K.G.H. A.G.T.G. assisted with mouse work. J.L.M. performed intrafemoral infections. M.G. performed analysis of the CBF-AML dataset. A.Z. performed the CIBERSORTx analysis. S.A. and A.S. performed eCLIP-seq. B.Y. analyzed eCLIP-seq data. K.G.H., A.G.T.G., N.B., O.I.G. and E.W. assisted with *in vitro* experiments. V.V., O.I.G., E.W., E.R.L. and J.E.D. edited the manuscript. J.A.K. and M.D.M coordinated AML patient consent, sample collection, and provided patient information. E.R.L. conceived the project, analyzed data and supervised the study. J.E.D. supervised research and secured funding for this study.

## Declaration of Interests

G.W.Y. is a cofounder, a member of the Board of Directors, on the Scientific Advisory Board, an equity holder and a paid consultant for Locanabio and Eclipse BioInnovations. G.W.Y. is a visiting professor at the National University of Singapore. G.W.Y.’s interests have been reviewed and approved by the University of California San Diego, in accordance with its conflict-of-interest policies. E.L.VN. is co-founder, member of the Board of Directors, on the SAB, equity holder, and paid consultant for Eclipse BioInnovations. E.L.VN.’s interests have been reviewed and approved by the University of California, San Diego in accordance with its conflict of interest policies. The authors declare no other competing interests.

## Tables

Table S1. Differentially Expressed Genes from RNA-seq of CD34^+^ HSPC Following miR-130a OE, TBL1XR1 KD and Kasumi-1 cells Following miR-130a KD (Excel Table)

Table S2. Differentially Expressed Proteins from Mass Spectrometry Analysis of CD34^+^ HSPC Following miR-130a OE (Excel Table)

Table S3. Quality Control Data for Chimeric AGO2 eCLIP

Table S4. miR-130a-Target Chimeras from AGO2 eCLIP in CD34^+^ CB and Kasumi-1 cells (Excel Table)

Table S5. CBF AML Patient Information (Excel table)

## STAR Methods

### RESOURCE AVAILABILITY

#### Lead Contact

Further information and requests for resources and reagents should be directed to and will be fulfilled by the lead contact, John Dick

#### Materials Availability

Recombinant DNA is listed in the Key Resources Table. The mass spectrometry data have been deposited to the ProteomeXchange Consortium (http://proteomecentral.proteomexchange.org) via the PRIDE partner repository with the dataset identifier PXD027331. Processed eCLIP-seq, RNA-seq and CUT&RUN data data have been deposited in NCBI’s Gene Expression Omnibus (GEO) under accession code (GSE181140). Raw RNA-seq and CUT&RUN data are under submission (EGA pending). This paper analyzes existing, publicly available data. These accession numbers for the datasets are listed in the Method details section.

Chimeric AGO2 eCLIP processing pipeline is available at (https://github.com/yeolab/eclip). Any additional information required to reanalyze the data reported in this paper is available from the lead contact upon request.

## Experimental Models and Subject Details 1.Primary Cells and Cell Lines

### 1.1 Samples Cryopreservation and Thawing

All samples were cryopreserved in 80% FBS (Multicell) and 20% DMSO (FisherScientific) solution added in 1:1 ratio. Samples were stored at −80°C for short-term storage or −150°C for long term storage. All primary cells and cell lines were thawed by slow, dropwise addition of thawing medium containing 50% X-VIVO 10 (Lonza), 50% FBS and DNaseI (100 ug/ml). Subsequently, cells were centrifuged at 1,450 rpm for 10 min at RT and resuspended in PBS (Multicell) + 2.5% FBS.

### 1.2. Cord blood processing and HSPC enrichment

Human cord blood (CB) samples were obtained with informed consent from Trillium Health Centre, Brampton Civic Hospital and Credit Valley Hospital, Ontario, Canada in accordance with guidelines approved by the University Health Network (UHN) Research Ethics Board. Cord blood samples were diluted 1:1 with PBS and mononuclear cells were enriched by density gradient centrifugation with lymphocyte separation medium (Wisent). Red blood cells were lysed with ammonium chloride solution (StemCell Technologies) and hematopoietic stem and progenitor cells were isolated with StemSep Human Hematopoietic Progenitor Enrichment Kit and Anti-Human CD41 TAC (StemCell Technologies) by negative selection according to manufacturer’s instructions. Lineage depleted (Lin-) CB cells were resuspended in PBS + 2% FBS and cryopreserved at −150°C.

### 1.3. Cell culture

CB cells were cultured in low cytokine conditions using X-VIVO 10 medium with 1% BSA, 1X penicillin-streptomycin (Gibco), 1X L-glutamine (Multicell) supplemented with the following cytokines: Flt3L (100 ng/ml), SCF (100 ng/ml), TPO (50 ng/ml) and IL-7 (10 ng/ml). Kasumi-1 and Flag-AML1-ETO Kasumi-1 cells were cultured in RPMI medium + 20% FBS, 1% Pen/Strep. Flag-AML1-ETO Kasumi-1 cell line was generously provided by S.W. Hiebert’s lab. Primary t(8;21) CD34^+^ blasts from AML patient samples were cultured in X-VIVO 10 medium supplemented with 20% BIT9500 (StemCell Technologies) and the following cytokines: IL-3 (10 ng/mL), SCF (50 ng/mL), IL-6 (10 ng/mL), Flt3L (50 ng/mL), G-CSF (10 ng/mL) and TPO (25 ng/mL) on MS5 stroma expressing human soluble CSF1. MS5 stroma cell line expressing human CSF1 was generously provided by M.D. Minden’s lab. Details of AML patient samples are outlined in Table S5. MS5-CSF1 stroma cells were grown in 6-well tissue culture treated plates seeded at a density of 1-1.5×10^5 MS5 stroma cells/well. MS5-CSF1 stroma cells and HEK293T cells were cultured in IMEM medium +10% FBS, 1% Pen/Strep. Cells were propagated in T75 tissue culture treated flasks and passaged every 2-4 days for up to 7-8 passages.

## 2. Xenotransplantation

### 2.1 Mouse Experiments

All mouse experiments were performed in accordance with the institutional guidelines approved by the University Health Network Animal Care Committee. All xenotransplantation experiments were performed with 8-12 week-old male and female NSG (NOD.Cg-*Prkdc^scid^ Il2rg^tm1Wjl^/Szj)* and NSG-SGM3 (NOD.Cg-*Prkdc^scid^* Il2rg^tm1Wj^ Tg(CMV-IL3, CSF2, KITLG)1Eav/MloySzj) (JAX) mice. Mice were sublethally irradiated (225 cGy) 24-48 hours before transplantation. Intrafemoral injections were performed on anesthetized mice by injecting 30 ul of cells resuspended in PBS into the right femur.

## Method Details

## 1. Lentiviral Vector Constructs, Lentiviral Transduction and Cell Sorting

### 1.1 Lentiviral Vectors

Lentiviral vectors for ectopic miRNA expression and stable knockdown have been described previously (Amendola et al., 2009; Gentner et al., 2009; Lechman et al., 2012, 2016). Briefly, pre-miRNA sequences were PCR amplified using sequence specific primers with Fse1 and Xho1 restriction enzyme sites (Key Resources Table) and genomic DNA from CD34^+^ CB cells. PCR product was subsequently purified using MinElute PCR Purification kit and digested with Fse1 and Xho1. Digested DNA fragments encoding human pre-miRNA sequences (∼350 nt in length) were cloned into the intron of LV.SFFV.intron.mOrange2 vector backbone. Empty vector backbone was used as an mO^+^ control (Ctrl). For stable knockdown of miR-130a, DNA fragment containing 8 tandem copies of imperfectly complementary sequence to mature miR-130a and Pac1 and Sph1 restriction enzyme sites (Key Resources Table) was ordered through GeneArt Gene Synthesis (ThermoFisher). Following restriction enzyme digest and gel electrophoresis, the DNA fragment was cloned into the LV.SFFV.GFP backbone using Rapid Ligation Kit (Thermo Fisher. Empty vector backbone was used as a negative control (Ctrl).

shRNA sequences for TBL1XR1 shRNA vectors were predicted using Sherwood algorithm (Knott et al., 2014) and ordered as Ultramer DNA oligos (IDT). The negative control for knockdown lentivirus (shCtrl) contains a sequence directed against Renilla luciferase in the pLBC2 backbone with BFP fluorescent reporter gene. Four different shRNA sequences for TBL1XR1 were PCR amplified using AmpliTaq Gold 360 Polymerase (ThermoFisher) with sequence specific primers containing BamH1 and Mlu1 restriction enzymes sites on the forward and reverse primers, respectively (Key Resources Table). PCR product was subsequently purified using MinElute PCR Purification kit (Qiagen) and digested with BamH1 and Mlu1. Digested PCR product and vector backbone were resolved by gel electrophoresis and extracted from the gel with Quick Gel Extraction Kit (Qiagen). Isolated PCR product was cloned into a Ultramir cassette (miR30) within a pRRL-based vector downstream of a SFFV promoter. Knock-down efficiency was assessed by qRT-PCR using TaqMan assay for TBL1XR1 (Thermo Fisher Cat#4331182; Hs00226564_m1) and GAPDH (Hs02758991_g1) and/or western blot assay using primer/probe sets and antibodies listed in the Key Resources Table.

### 1.2 Lentivirus Production and Titration

VSV-G pseudotyped lentiviral vector particles were produced in 293T cells using 2^nd^ and 3^rd^ generation packaging protocols described previously (Follenzi et al., 2000; Guenechea et al., 2000). 293T cells were seeded in 15 mm plates with approximately 9-10×10^6 cells per plate one day before transfection to achieve ∼80% confluency at the time of transfection. Subsequently, cells were transfected using CalPhos Mammalian Transfection Kit (Takara) according to the manufacturer’s protocol. Fresh AMEM medium + 1%FBS, 1X L-glutamine, 1X penicillin-streptomycin and sodium butyrate (1mM) was added to transfected plates ∼16 hours post-transfection. Virus-containing medium was collected 48 hours post-transfection and cell debris was pelleted by centrifugation at 1,800 rpm for 15 min at RT. Subsequently, supernatant was filtered using 0.45 µm filter, virus particles were concentrated using ultracentrifugation at 22,000 rpm for 2.5 hr at 4°C and stored at −80°C. Lentiviruses were titrated on 293T cells.

### 1.3 Lentiviral Transduction

Transduction of Lin-CB CD34^+^CD38^-^ cells and other HSPC populations was performed in low cytokine conditions described above. Sorted CD34^+^CD38^-^ CB cells were transduced with lentiviruses at a density of 5×10^5 cell/ml. Fresh medium was added 24 hours post-transduction and cells were harvested 72 hour post-transduction and subsequently transplanted into mice or collected for other experimental procedures. Transduction efficiency was determined by removing 10% of the transduced cells and measuring the percentage of mOrange^+^, GFP^+^ or BFP^+^ cells by flow cytometry.

Transduction of Kasumi-1 and Flag-AE Kasumi-1 cells was performed at a density of 2×10^6 cells/ml in RPMI medium with 20% FBS, 1% Pen/Strep. Cells were infected in 24-well suspension plates and centrifuged at 1,400 rpm for 60 min at 32°C and fresh medium was added on the next day. For xenotransplantation of primary CD34^+^ t(8;21) AML blasts, cells were enriched using human CD34 Microbead Kit (Miltenyi Biotec) according to manufacturer’s instructions. Enriched CD34^+^ blasts were transduced at a density of 1.5-9×10^6 cells/ml in X-VIVO + 20% BIT 9500 supplemented with the following cytokines IL-3 (10 ng/ml), SCF (50 ng/mL), IL-6 (10 ng/mL), Flt3L (50 ng/mL), G-CSF (10 ng/mL) and TPO (25 ng/mL) and fresh media was added on the following day. Transduction efficiency was determined 72 hours post-infection by flow cytometry. For AML 120567, 3×10^5 CD34^+^ blasts were transplanted per mouse. For AML 130199, 1×10^6 CD34^+^ blasts were transplanted per mouse. For in vitro experiments, CD34^+^ blasts were transduced as described and transferred to MS5 CSF1 stroma 24 hours post-transduction. Blasts were sorted for GFP^+^ cells 72 hours post-transduction, cultured for additional 5 days and then analyzed for cell surface marker expression by flow cytometry.

## 2. Cell Sorting and Flow Cytometric Analysis

Cord blood cells, primary AML cells and cell lines were resuspended at 10^6^ cells/ml in PBS + 2% FBS for surface antibody staining and antibodies were used as indicated (Key Resources Table). Cells were stained for 20 min at room temperature and washed with PBS. Cell sorting was performed at FACSAria III (BD Biosciences), SH800 (Sony) or MoFlo (Beckman Coulter) sorters or analyzed on Canto II or Celesta instruments (BD). Antibodies are listed in the Key Resource Table.

## 3. Mouse Experiments

### 3.1 Xenotransplantation

For CB transplantation, progeny of ∼1×10^4 CD34^+^CD38^-^ CB cells were injected per mouse 72 hours post-transduction. The miRNA OE screen was performed with 3 independent cord blood pools each transduced with 3 different miRNA OE and 1 control lentivirus and transplanted into 4-5 mice per experimental condition. For miR-130a OE repopulation assays, 2-4 independent CB pools were transduced for each experiment and transplanted into 8-10 mice per experimental condition. Mice were sacrificed after 12 or 24 weeks of repopulation and right femur (RF) and bone marrow (left femur and tibia) were flushed with 1ml of PBS + 2.5% FBS. Spleens were weighted, crushed and collected using cell strainer caps. All cells are centrifuged at 1,450 rpm for 10 min at RT and resuspended in 500 ul of PBS + 2.5% FBS. Cells were counted in ammonium chloride using Vicell XR (Beckman Coulter) and used for flow cytometric analysis or other assays.

Secondary transplantations for miR-130a OE studies and TBL1XR1 KD studies were performed by combining cells from 2-6 primary recipient mice and depleting murine cells using the Mouse Cell Depletion Kit (Miltenyi Biotec) according to manufacturer’s instructions. Murine-depleted cells were sorted for hCD45^+^mO^+^ or hCD45^+^BFP^+^ cells and transplanted at defined doses into NSG or NSG-SGM3 mice using intrafemoral injections. Secondary recipient mice were sacrificed at week 8-12, right and left femur were flushed and analyzed by flow cytometry. Engraftment was scored positive if the percentage of hCD45^+^ BFP^+^/mO^+^ cells was >0.05% in the bone marrow. LT-HSC frequency was estimated by linear regression analysis and Poisson statistics using the publicly available online software ELDA (Key Resource Table).

### 3.2 Lin-Cell Depletions or CD34^+^ Enrichment

For flow cytometric analysis of HSPC populations from miR-130a OE and miR-130a KD xenografted mice, cells RF and BM of 2-4 mice were enriched for human HSPC with Mouse/Human Chimera Enrichment Kit (STEMCELL Technologies), StemSep Human Hematopoietic Progenitor Enrichment Kit (STEMCELL Technologies) and Anti-Human CD41 TAC (STEMCELL Technologies, 14050). For enrichment of CD34^+^ blasts from peripheral blood of CBF-AML primary patient samples, Direct CD34^+^ Progenitor Isolation Kit (Miltenyi Biotec) was used according to manufacturer’s instructions. For secondary transplantation experiments and analysis of HSPC from TBL1XR1 KD xenografts, Mouse Cell Depletion Kit (Miltenyi Biotec) was used to deplete murine cells and isolated human cells were subsequently analyzed by flow cytometry or sorted using FACS for secondary transplantations or RNA-seq. Cells from 2-4 individual mice were combined prior to depletion of murine cells.

### 3.3 Flow Cytometric Analysis of Xenografts

Cells from RF, BM, or spleen of xenografted mice (50 ul) were stained in 4 ml polystyrene tubes (Falcon, 352058) or 96 well U bottom plates (Corning, 351177) for 30 min at 4°C. Following antibodies (Key Resource Table) were used: 1. For the Pan Lin stain, CD3 FITC (1:100), GlyA PE Cy5 (1:100) or GlyA PE (1:100), CD41 PE Cy7 or CD41 PE Cy5 (1:100), CD33 APC, CD45 APC Cy7 (1:100) or CD45 V500(1:50), CD19 V450 (1:100) or CD19 PE C7 (1:100); 2. For the erythroid stain, CD71 FITC (1:100), GlyA PE Cy5 (1:100), CD117 PE cy7 (1:100), CD36 APC (1:50), CD34 APC Cy7 (1:200); 3. For the myeloid stain, CD45 FITC (1:100), CD49d PE Cy5 (1:50), CD14 PE Cy7 (1:100), CD66b APC (1:100), CD16 APC Cy7 (1:50), CD33 V450 (1:100). For the full stem and progenitor hierarchy stain, murine-depleted cells from TBL1XR1 KD and Ctrl xenografts were stained with the following antibodies: CD45RA FITC (1:50), CD45 V500 (1:50), Flt3 BV711 (1:50), CD7/CD10/CD19 AF700 (1:100), CD38 PE-Cy7 (1:100), CD90 APC (1:50), CD34 APC Cy7 (1:200) and CD49f PE-Cy5. For the miR-130a OE and KD, xenografts depleted of murine and lineage committed cells were stained with the following antibodies: Flt3-biotin (1:50), STV-FITC (1:100), CD7/10 PE-Cy5 (1:100), CD38 PE-Cy7 (1:100), CD90 APC (1:50), CD34 APC Cy7 (1:200), CD45RA BV421 or CD45RA FITC (1:100).

## 4. RNA-seq

### 4.1 RNA isolation

Sorted populations from cord blood or Kasumi-1 cells (∼5×10^3 - 5×10^4 cells) were washed in PBS and centrifuged at 1,450 rpm for 10 minutes and RT. Cell pellets were frozen at −80°C and total RNA was extracted using RNeasy Plus Micro Kit (Qiagen) or PicoPure RNA Isolation Kit (ThermoFisher) according to manufacturer’s instructions. RNA integrity was assessed using Agilent Bioanalyzer (RNA Pico chip Agilent Technologies) and next-generation sequencing libraries were prepared at the Center of Applied Genomics, SickKids Hospital. SMART-Seq V4 Ultra Low Input RNA kit for sequencing (Clontech) was used to generate the cDNA samples. Subsequently, next-generation sequencing libraries were prepared with Nextera XT DNA Library Preparation Kit (Illumina). Equimolar quantities of libraries were pooled and sequenced on the Illumina HiSeq 2500 platform generating 125-bp paired-end sequencing reads to achieve depth of approximately 40-50M reads per sample. All RNAseq experiments were performed *in vitro* in triplicates using three individually transduced samples.

### 4.2 RNA-seq Processing and Pathway Enrichment Analysis

Raw sequencing reads were trimmed to remove the adaptors and aligned against hg38 human genome with STAR v2.5.2b (Dobin et al., 2013) using GENCODE (Frankish et al., 2019). Read counts were generated using HTSeq counts v0.7.2 (Anders et al., 2015) and general statistics were obtained from picard/2.6.0 using Default parameters. Subsequently, unstranded counts were normalized using normalization factor and annotated using BioMart_ENSG_hg38. For TBL1XR1 KD in CB, STAR v2.7.2 and HTSeq was used to count reads over GENCODE v32 features using default parameters. EdgeR_3.28.1 (Robinson et al., 2010) was used to fit a glmQ model to generate estimation of differential expression between miR-130a OE in CB and miR-130a KD in Kasumi and TBL1XR1 KD in CB conditions and their respective control samples. A score was calculated using the formula sign(logFC)*-log10(p value) to rank all genes from top upregulated to downregulated and was subsequently used in pathway enrichment analysis.

#### 4.2.1 Pathway Enrichment Analysis of miR-130a OE in CB

The ranked gene lists were used in GSEA_4.1.0 using the Baderlab gene set file containing pathways from multiple databases (June 01 2020, http://baderlab.org/GeneSets/), using as parameters 2000 permutations and gene set size between 10 and 500. EnrichmentMap 3.3.2 in Cytoscape 3.8.2 was used to visualize the GSEA enrichment results and AutoAnnotate 1.3.3 was used to cluster and label functional modules. Significant overlap at a p value of 0.05 between the downregulated DE proteins between miR-130a OE and Control and DIANA-TarBase v7.0 miR-130a targets targets was visualized in Cytoscape/EnrichmentMap using the post-analysis function. GSEA was also run on the following selected gene sets. Cell cycle gene sets used in the miR-130a OE GSEA including G1 S DNA damage, Myc targets and positive regulation of mitotic cell cycle were obtained from the GO biological process and MSig C2 databases. Proliferative and quiescent gene sets were obtained from Forsberg et al. comparing gene expression of quiescent and cytokine-induced mobilized HSC in mouse (Forsberg et al., 2010). RA-target genes were extracted from several publicly available data (Balmer and Blomhoff, 2002; Cabezas-Wallscheid et al., 2017; Ghiaur et al., 2013). For the RA-target gene dataset from Ghiaur et al. 2013, DE genes between CD34^+^CD38^-^ and CD34^+^CD38^+^ cells from bone marrow donors and predicted RA targets (Table S1) were used. RA-target genes from Balmer and Blomhoff 2002 dataset were extracted, genes were converted to human orthologs using g:convert (https://biit.cs.ut.ee/gprofiler/convert) and separated into upregulated and downregulated by RA.

#### 4.2.2 Pathway Enrichment Analysis of miR-130a KD in Kasumi

The ranked gene lists were used in GSEA_4.1.0 using the Baderlab gene set file containing pathways from multiple databases (June 01 2020, http://baderlab.org/GeneSets/), using as parameters 2000 permutations and gene set size between 10 and 500. EnrichmentMap 3.3.2 in Cytoscape 3.8.2 was used to visualize the GSEA enrichment results and AutoAnnotate 1.3.3 was used to cluster and label functional modules. Overlap between the enriched pathways and DIANA-TarBase v7.0 miR-130a targets was assessed using Fisher’s exact test and significant overlap at p value 0.05 was visualized using the post-analysis function.

#### 4.2.3 Pathway Enrichment Analysis of TBL1XR1 KD in CB

The ranked gene lists were used in GSEA_4.1.0 using the Baderlab gene set file containing pathways from multiple databases (June 01 2020, http://baderlab.org/GeneSets/), using as parameters 2000 permutations and gene set size between 10 and 500. EnrichmentMap 3.3.2 in Cytoscape 3.8.2 was used to visualize the GSEA enrichment results and AutoAnnotate 1.3.3 was used to cluster and label functional modules. Overlap between the enriched pathways and eCLIP miR-130a targets was assessed using Fisher’s exact test and significant overlap at p value 0.05 was visualized using the post-analysis function. The same gene sets used in the miR-130a OE analysis were used in the GSEA of TBL1XR1 KD DE genes including the proliferative and quiescent gene sets (Forsberg et al., 2010) and Myc targets and positive regulation of mitotic cell cycle obtained from the GO biological process and MSig C2 databases. RA-upregulated and downregulated genes were obtained from Cabezas-Wallscheid et al. 2017 comparing in vitro ATRA treatment of murine HSC to untreated HSC (LSK CD150+CD48−CD34−) and RA-target genes from Balmer and Blomhoff (2002).

## 5. Mass Spectrometry and Downstream Analysis of miR-130a Targets

### 5.1 Label free-mass spectrometry samples preparation

Human Lin-CB cells from 3 independent cord blood pools were transduced individually with miR-130a OE and control lentiviruses and collected for FACS 72 hours post-transduction. Equal number of sorted CD34^+^mO^+^ cells (1×10^5) were washed twice with ice-cold PBS and resulting samples were subjected to sample preparation similar to Schoof et al., 2016 (Schoof et al., 2016). Cells were lysed using 20 ul of lysis buffer (6M Guanidinium Hydrochloride, 10 mM TCEP, 40 mM CAA, 100 mM Tris pH8.5). Samples were boiled for 5 minutes 95°C, after which they were sonicated on high for 3×10 seconds in a Bioruptor sonication water bath (Diagenode) at 4°C. Samples were diluted 1:3 with 10% Acetonitrile, 25 mM Tris pH 8.5, LysC (MS grade, Wako) was added in a 1:50 (enzyme to protein) ratio, and samples were incubated at 37°C for 4hrs. Samples were further diluted to 1:10 with 10% Acetonitrile, 25 mM Tris pH 8.5, trypsin (MS grade, Promega) was added in a 1:100 (enzyme to protein) ratio and samples were incubated overnight at 37°C. Enzyme activity was quenched by adding 2% trifluoroacetic acid (TFA) to a final concentration of 1%. Prior to mass spectrometry analysis, the peptides were desalted and fractionated on in-house packed SCX Stagetips. For each sample, 3 discs of SCX material (3M Empore) were packed in a 200ul tip, and the SCX material activated with 80 μl of 100% Acetonitrile (HPLC grade, Sigma). The tips were equilibrated with 80 μl of 0.2% TFA, after which the samples were loaded using centrifugation at 4,000x rpm. After washing the tips twice with 100 μl of 0.2% TFA, two initial fractions were eluted into clean 500 μl Eppendorf tubes using 75 mM and 300 mM ammonium acetate in 20% Acetonitrile, 0.5% formic acid respectively. The final fraction was eluted using 5% ammonium hydroxide, 80% Acetonitrile. The eluted peptides were frozen on dry ice and concentrated in an Eppendorf Speedvac, and re-constituted in 1% TFA, 2% Acetonitrile for Mass Spectrometry (MS) analysis.

### 5.2 Mass Spectrometry Acquisition

For each sample, peptides were loaded onto a 2cm C18 trap column (ThermoFisher 164705), connected in-line to a 50 cm C18 reverse-phase analytical column (Thermo EasySpray ES803) using 100% Buffer A (0.1% Formic acid in water) at 750 bar, using the Thermo EasyLC 1000 HPLC system in a single-column setup and the column oven operating at 45°C. Peptides were eluted over a 200 minute gradient ranging from 5 to 48% of 100% acetonitrile, 0.1% formic acid at 250 nl/min, and the Orbitrap Fusion (Thermo Fisher Scientific) was run in a 3 second MS-OT, ddMS2-IT-HCD top speed method. Full MS spectra were collected at a resolution of 120,000, with an AGC target of 4×10^5^ or maximum injection time of 50ms and a scan range of 400–1500 m/z. Ions were isolated in a 1.6m/z window, with an AGC target of 1×10^4^ or maximum injection time of 35ms, fragmented with a normalized collision energy of 30 and the resulting MS2 spectra were obtained in the ion trap. Dynamic exclusion was set to 60 seconds, and ions with a charge state <2, >7 or unknown were excluded. MS performance was verified for consistency by running complex cell lysate quality control standards, and chromatography was monitored to check for reproducibility. The mass spectrometry data have been deposited to the ProteomeXchange Consortium (http://proteomecentral.proteomexchange.org) via the PRIDE partner repository with the dataset identifier PXD027331.

### 5.3 Label-free Quantitative Proteomics Analysis

The raw files were analyzed using MaxQuant version 1.5.2.8 (Cox and Mann, 2008) and standard settings. Briefly, label-free quantitation (LFQ) was enabled with a requirement of 2 unique peptides per protein, and iBAQ quantitation was also enabled during the search. Variable modifications were set as Oxidation (M), Acetyl (protein N-term), Gln->pyro-Glu and Glu->pyro-Glu. Fixed modifications were set as Carbamidomethyl (C), false discovery rate was set to 1% and “match between runs” was enabled. The resulting protein groups file was processed with an in-house developed tool (PINT) (Wojtowicz et al., 2016), which imputes missing LFQ values with adjusted iBAQ values. Briefly, the distributions of iBAQ intensities for each sample are adjusted to overlap with the LFQ intensity distributions using median-based adjustment, enabling the direct imputation of missing LFQ values with adjusted iBAQ values for those proteins that did not have LFQ values across all the samples. Final list of 6735 proteins identified and quantified in all samples was generated by simultaneously filtering for reverse hits, contaminants and only those proteins observed in 3 biological replicates (n=3) in at least one group. Significance of protein ratios between Control and miR-130a OE samples were estimated for each biological repeat and subjected to a statistical analysis in R using limma_3.42.2 with Benjamini-Hochberg adjustment. The 6735 detected proteins were ranked from top upregulated to downregulated in miR-130a OE compared to control using the t statistics values from the limma paired t-test and this rank file was subsequently used in the pathway analysis (Table S2).

### 5.4 Pathway Analysis and the Enrichment Map Visualization

GSEA_4.10 was performed using the ranked list from the proteomics data and Baderlab gene set file containing pathways from multiple databases (June 1, 2020, http://baderlab.org/GeneSets/) using as parameters of 2000 permutations and gene set size between 10 and 500. EnrichmentMap 3.3.2/ Cytoscape 3.8.2 was used to visualize 145 and 128 gene sets that were significantly enriched at p value 0.05 in genes upregulated and downregulated in miR-130a OE, respectively. To identify direct miR-130a targets, a proteomics analysis was combined with computationally predicted and experimentally validated miR-130a targets from the mirDIP v1.0 (microRNA data integration portal) and DIANA-TarBase v7.0 databases, respectively. To estimate the enrichment of miR-130a targets in downregulated proteins, a Wilcoxon rank sum test was applied between the ranked list of DE proteins between miR-130a OE and control and Tarbase predicted targets. The specificity of the enrichment of miR-130a targets in downregulated proteins was confirmed by comparing it to the ones obtained for the other 6 other members of the miR-130 family, 7 randomly chosen miRNAs, and 2000 random gene lists by plotting the observed and random p-values on a histogram in R. GSEA was also used to confirm this enrichment by using a gene set of all Tarbase miR130a targets present in more than 2 independent studies. Significant overlap at a p value of 0.05 between the downregulated DE proteins between miR-130a OE and Control and Tarbase predicted targets was visualized in Cytoscape/EnrichmentMap using the post-analysis function. Furthermore, 9 miR-130a targets overlapping between both mirDIP and Tarbase databases with downregulated proteins at p value<0.05 were further visualized using the heatmap.2 function in R and Cytoscape/EnrichmentMap. GSEA was also used to confirm the enrichment of the 9 targets in downregulated proteins.

## 6. Chimeric AGO2 eCLIP-seq and Post-Analysis with miR-130a-target chimeras

### 6.1 Chimeric AGO2 eCLIP

Chimeric eCLIP for miRNA-mRNA chimeras (Van Nostrand EL, Shishkin AA, & Yeo GW, in preparation) was performed to identify global miRNA-target interactions in CD34^+^ CB and Kasumi-1 cells. Briefly, the chimeric eCLIP method utilizes the enhanced crosslinking immunoprecipitation protocol (Van Nostrand et al., 2016) with the following changes: after initial washes of AGO2 immunoprecipitates, samples were treated with T4 PNK 3’ phosphatase minus (NEB) to phosphorylate RNA fragment 5’ ends, followed by T4 RNA ligation without adapter to encourage chimeric ligation products. After standard eCLIP adapter ligation and gel electrophoresis, reverse transcription was performed under modified Mn^2+^ buffer conditions (Van Nostrand et al., 2017) to generate cDNA libraries, and PCR amplified as per the eCLIP protocol. To generate mir130-targeted AGO2 eCLIP libraries, the content of miR130a chimeras in 0.25 µl of cDNA library was quantified by qPCR (NEB LUNA Universal qPCR 2x Master Mix) using primer q7c and primer mir130a-enrich (Key Resource Table). The remainder of the libraries were amplified (NEB Q5 High-Fidelity 2x Master Mix) with primer Q23C and primer mir130a-enrich, using the following cycle parameters: 98°C, 30s; (98°C, 15s; 65°C, 30s; 72°C, 40s) × 6 cycles; (98°C, 15s; 72°C, 45s) × 7 cycles; 72°C, 1 min. PCR products were purified (AMPure XP beads, Beckman Coulter; 1.8 × bead to sample ratio), and amplified in a second PCR reaction using standard Illumina D5X and D7X index primers to generate sequencing libraries, using the following cycle parameters: 98°C, 30s; (98°C, 15s; 72°C, 1 min) × 12 cycles; 72°C, 1 min. PCR products were purified (AMPure XP beads; 1.5 × bead to sample ratio). All libraries were sequenced in PE100 mode on the Illumina NovaSeq-6000 platform.

### 6.2 Chimeric AGO2 eCLIP data analysis

Processing and bioinformatics analysis was based on previously described methods (Moore et al., 2015), with the following modifications: Ago2 peaks were assigned using the eCLIP processing pipeline available at (https://github.com/yeolab/eclip). Briefly, unique molecular identifiers (UMIs) were assigned to each read using umi_tools (1.0.0) and trimmed of adapters with cutadapt (1.14.0). Reads were then mapped to repeat elements (RepBase release 18.05) using STAR (2.7.6a). Reads that did not align to repeat elements were then aligned to hg19 with STAR and UMI-collapsed with umi_tools. These PCR-deduped reads were then used to identify local enrichments with CLIPper (3.0) and normalized above SMInput using custom scripts overlap_peakfi_with_bam.pl and compress_l2foldenrpeakfi_for_replicate_overlapping_bedformat.pl.

To identify chimeras within total eCLIP datasets, UMI-tagged reads were adapter trimmed, sorted and collapsed, leaving only unique sequences. These sequences were then indexed with Bowtie2 (2.2.6) and used to reverse-map miR sequences (miRBase release 22.1). Each miR-mapped read was filtered to select one miR per unique sequence, prioritizing positive stranded reads and minimizing mismatches or indels. To identify chimeras, miR-mapped reads were then uncollapsed and those which contained an mRNA portion of at least 18nt were mapped to hg19 with STAR (2.7.6a). Chimeric reads were then PCR-deduplicated and overlapped with identified Ago2 peaks.

Targeted miR-130a-specific eCLIP reads were UMI-tagged and adapter trimmed in similar fashion as what was done with total datasets. Chimeras from these targeted libraries were identified as hg19-mapped reads that contain expected primer sequences. These reads were then PCR-deduplicated and used to identify enriched regions with Clipper. For both total and targeted chimeric datasets, peaks were annotated (Gencode v19) to identify the fractions of bound genic regions (ie. CDS, UTR, intron). De-novo motif analysis using HOMER (4.9.1-6) was also performed to ensure that the most enriched motif corresponds to the expected miR seed. Sequencing and genome mapping statistics are listed in Table S3.

### 6.3 Pathway Analysis of miR-130a-target chimeras in CB and Kasumi-1 cells

Pathway enrichment analysis was performed with the web based tool g:Profiler (https://biit.cs.ut.ee/gprofiler/gost) using the Baderlab gene set file (June 1, 2020, http://baderlab.org/GeneSets/<u>) and</u> visualized using EnrichmentMap 3.3.2/ Cytoscape 3.8.2 and AutoAnnotate 1.3.3. Significant overlap at a p value of 0.05 between the downregulated DE proteins between miR-130a OE and Control and the list of miR-130a-target chimeras was calculated using a Wilcoxon rank sum test and visualized in Cytoscape/EnrichmentMap using the post-analysis function. The same analysis was applied to DE transcripts between miR-130a KD and Control and the list of miR-130a-target chimeras from Kasumi-1 cells.

## 7. Cell Cycle Analysis

Cell cycle analysis was performed with the Click-iT EdU AF647 Flow Cytometry Assay kit according to manufacturer’s instructions. Lin-CB was used to sort LT-HSC, ST-HSC and CD34+CD38+ progenitors using the following antibodies (BD): CD34 APC-Cy7 (1:200), CD38 PE-Cy7 (1:100), CD90 APC (1:50), CD45RA FITC (1:50) and CD49f PE-Cy5 (1:50) and PI. Sorted subpopulations (approximately 7×10^3^-1×10^4^) were cultured in low cytokine conditions overnight and transduced with miR-130a OE or Control viruses 24 hours later. EdU was added into the medium at 10 uM concentration and cells were pulsed for 1.5-2 hr. Subsequently, cells were fixed and permeabilized as described in the protocol. Cells were stained with Ki67-FITC antibody (1:30, BD) in PermWash solution overnight at 4°C. The next day, cells were stained with DAPI (1:5000) to label DNA and analyzed by flow cytometry.

## 8. miR-130a expression in AML

### 8.1 Quantitative RT-PCR for expression levels of miR-130a in CBF AML

Peripheral blood samples from t(8;21) and inv(16) patients (Table S5) were thawed as described and CD34^+^ blasts were enriched with Direct CD34^+^ Progenitor Isolation Kit (Miltenyi) or sorted by FACS. Following column enrichment, RNA was extracted from approximately 5×10^5 CD34^+^ or CD34^-^ cells with MirVana miRNA Isolation Kit (ThermoFisher) and RNA was reverse transcribed with TaqMan MicroRNA reverse transcription kit using miR-130a-3p (Thermo Fisher Cat#4427975; ID 000454) and RNU48 (ID001006) specific primers purchased from TaqMan Small RNA assay (ThermoFisher). Subsequently, qPCR was performed on the ABI-SDS7900HT instrument using the corresponding Taqman primer/probe sets. Relative quantification was performed using the ΔΔCt method normalized to the levels of RNU48 and miR-130a expression in PBMC from 3 healthy volunteers was used as a control. The following AML cell lines were used to measure the expression level of miR-130a: OCI-AML3, OCI-AML2, MOLM13, H260, NB-4, Kasumi-1, Flag-AE Kasumi-1 and U937. Cells (∼5×10^5) were collected in triplicates, washed in PBS and centrifuged at 1,450 rpm for 10 minutes at 4°C. RNA was extracted with RNeasy Plus Mini Kit (Qiagen), RNA integrity was assessed using Agilent Bioanalyzer (RNA Pico chip Agilent Technologies) and 10 ng of RNA was used in the RT reaction. RT-qPCR was performed using the TaqMan Small RNA Assays as described above. Expression levels of miR-130a in AML cell lines were represented relative to the expression levels in Kasumi-1 cells.

### 8.2 miRNA microarray in CBF AML patient samples

RNA was extracted from peripheral blood cells (1×10^6-1×10^7) using the RNeasy Mini kit (Qiagen). Briefly, samples were lysed, and total RNA was collected by column extraction according to the manufacturer’s instructions. Samples were then examined by Nanostring, a probe-based assay that detects 827 common miRNAs (https://www.nanostring.com/products/ncounter-assays-panels/immunology/mirna/). The output of the assay was analyzed by nSolver 4.0 where the mean of the negative spike-in control was used as the threshold of microRNA detection. With the expression profile table generated, patients were then stratified by miR-130a expression using a median split and a Kaplan-Meier curve was drawn by GraphPad Prism 7.

### 8.3 TCGA analysis of CBF AML patients

Publicly available data from the TCGA-LAML database for which miRNA expression data was available was used to study the miR-130a expression pattern. TMM normalized count per million counts (CPM) were clustered using the heatmap.2 function available from the gplots R package (gplots_3.1.1). The 4 main AML clusters separating the samples in very high, medium, low and very low miR-130a expression groups were retrieved from the heatmap cluster results using the R stats cutree function. A boxplot was constructed using CPM values for the 4 AML groups. One Way Analysis of Variance (ANOVA) was applied to the data to test the significance of the differences in the mean values between the groups. TCGA-LAML bulk RNAseq dataset was then used to determine the association of miR-130a expression with different AML subtypes. Total of 188 AML patients including cytogenetically normal (168) and complex AML (13 with inv(16) and 7 with t(8;21)) were included in the analysis. Expression levels of miR-130a in inv(16) and t(8;21) were compared to the other subtypes using normalized log2 CPM counts. LSC17 score was calculated on RPKM data of each TCGA-LAML patient sample data using regression coefficients of the 17 signature genes from Ng. et al. (Ng et al., 2016). A scatter and a boxplot plot were used to visualize the association of LSC17 score with miR-130a expression and clusters respectively. ANOVA was applied to the data to test the significance of the differences in the mean values between the 4 clusters.

### 8.4 GSVA and CIBERSORTx Analysis and deconvolution of AML signatures from scRNAseq

Deconvoluted AML data was obtained from Zeng et al (Zeng et al., submitted). Briefly, a signature matrix was generated using scRNA-seq data from van Galen et al. (van Galen et al., 2019) using expression profiles of seven malignant cell types: Leukemia Stem and Progenitor Cell (LSPC)-Quiescent, LSPC-Primed, LSPC-Cycle, Granulocyte-Monocyte-Progenitor (GMP)-like, ProMono-like, Mono-like, conventional Dendritic Cell (cDC)-like, as well as seven non-malignant cell types: T, Cytotoxic T Lymphocyte (CTL), Natural Killer (NK), B, Plasma, Monocyte, and cDC. LSPC populations were re-annotated in Zeng et al. Deconvolution using this signature matrix performed using CIBERSORTx with S-mode batch correction in absolute mode, and applied on TPM-normalized gene expression data from 173 TCGA AML patients and Kasumi-1 cells. Deconvolution analysis was also performed using RNAseq data from Kasumi-1 transduced with miR-130a KD or lentivirus. Inferred abundances of the seven malignant cell types were normalized to a total of 1, such that the score for a given population represents the proportion of total leukemic cells belonging to that population. For TCGA analysis, principal component analysis was performed based on malignant cell composition and clusters were labeled based on Zeng et al. Single sample GSEA was run on bulk TCGA-LAML RNAseq RPKM data by using the gsva function available from GSVA_1.34.0 using tumor-derived HSC/progenitor-like, GMP and myeloid signature gene-sets derived from)/σ (where μ is the mean of all x values and σ is the standard deviation of all x values). Each sample was classified as highly enriched in HSC/progenitor-like, GMP or myeloid gene-sets when the gsva score was greater than 1 standard deviation. A boxplot was constructed by plotting the miR-130a expression values for each tumor derived signature classified as highly or lowly enriched samples. A Student’s t test was applied to test the significance of the differences in the mean values between each 2 groups.

## 9. Western blotting and Immunoprecipitation of Flag-tagged AML1-ETO

### 9.1 Western blot assay

Sorted CD34^+^CD38^-^ cord blood cells or Kasumi-1 cells (∼5×10^4 - 2×10^5) were centrifuged at 1450 rpm for 10 min at RT and washed with PBS. Washed cell pellets were lysed in RIPA buffer (ThermoFisher) containing protease and phosphatase inhibitors (ThermoFisher). Subsequently, samples were centrifuged at 16,000xg for 15 min at 4°C and supernatants were utilized for Western blot assay. Western blot assay was performed on the automated Simple Western capillary platform (Wes, ProteinSimple) using 12-230 kDa or 66-440 kDa capillary cartridges according to manufacturer’s protocol. All antibodies listed in the Key Resources Table were titrated prior to use on lysates from CD34^+^ cord blood cells or Kasumi-1 cells and subsequently used at the listed dilutions.

### 9.2 Immunoprecipitation of Flag-AML1-ETO

Immunoprecipitation experiments were performed in Flag-AE Kasumi-1 cell line which was generously provided by Scott Hiebert’s lab. Flag-tagged AML1-ETO Kasumi-1 cells were transduced with miR-130a KD and control lentiviruses as described and sorted for GFP^+^ cells 72 hours post-transduction. Sorted cells (7.5×10^5-1.5×10^6) were centrifuged at 1,450 rpm for 10 min at RT and washed with PBS. Cell pellets were frozen at −80C until IP experiments were performed. IP experiments were performed using EZview Red ANTI-FLAG M2 Affinity Gel (Sigma) according to manufacturer’s instructions. Briefly, 20 ul of gel suspension was used per reaction. Gel beads were washed two times with 500 ul of TBS buffer (50mM Tris HCl, 150 mM NaCl, pH 7.4) by centrifugation ar 8,200xg for 30 seconds at RT. Cells were lysed in 500 ul of lysis buffer (50 mM Tris HCl pH 7.4, 150 mM NaCl, 1 mM EDTA, 1% TRITON X-100 and 1X protease inhibitors). Cell lysates were sonicated with Sonic Dismembrator Model 500 (Fisher Scientific) using 10% amplitude and 3 cycles with 10 seconds on and 20 seconds off pulses. Subsequently, samples were centrifuged at 20,000xg for 20 minutes and 4°C to pellet the debris and supernatant was collected. A fraction of each lysate (50 ul) was saved as an input and the remainder was added to the washed beads. Samples were incubated by gentle rotation overnight at 4°C. Next, beads were centrifuged at 8,200xg for 30 seconds and supernatant was discarded. Beads were washed 3 times with 500 ul of TBS and Flag-AML1-ETO was eluted under native conditions by competition with 3XFLAG peptide. For elution, 50 ul of TBS containing 150 ng/ul of FLAG-peptide was added to the washed beads. Samples were incubated by gentle rotation for 30 minutes at 4°C, followed by centrifugation at 8,200xg for 30 seconds. Supernatant containing the eluted Flag-AML1-ETO complex was transferred to fresh tubes, stored at −20°C or used right away in western blot assay.

## 10. CUT&RUN assay and Bioinformatic Analysis

### 10.1 CUT&RUN Assay for Flag-AML1-ETO binding occupancy

Flag-AML1-ETO Kasumi-1 cells were sorted 72 hr post-transduction and subsequently 2×10^5 cells per condition were used in CUT&RUN assay. Cells were washed in PBS and CUT&RUN assay was performed as previously described (Skene and Henikoff, 2017; Skene et al., 2018). Protein A-Micrococcal nuclease (pA-MNase) fusion protein and yeast spike-in DNA were kindly provided by S. Henikoff’s lab. Briefly, cells were washed twice with a wash buffer and activated Concanavalin A Beads (Bangs Laboratories) were added dropwise while vortexing the samples. Wash buffer was removed by beads separation on a magnet and antibody buffer containing 0.0125% digitonin and Anti-Flag, anti-ETO or mouse IgG control antibody (Key Resources Table) were added to the beads. Samples were incubated on a rotator overnight at 4°C, followed by incubation with rabbit anti-mouse secondary antibody for 1 hour at RT. Addition and activation of pA-MNase and isolation of soluble DNA was performed as previously described (Skene et al., 2018). DNA was extracted with the MinElute PCR Purification kit (Qiagen) and DNA libraries were prepared with NEBNext Ultra II DNA Library Prep Kit for Illumina (NEB) and NEBNext Multiplex Oligos for Illumina (NEB) according to manufacturer’s instructions. The number of PCR cycles for library preparation was determined from qPCR based on the Ct value. For each sample, 1 ul of DNA, 5 ul of Power SYBR Green PCR Master Mix (ThermoFisher), 0.4 ul of each forward (10 uM) and reverse primer (10 uM) and 3.2 uL of H_2_O were combined and qPCR was performed according to manufacturer’s instruction. AMPure XP beads (Beckman) were used for cleanup of PCR reactions as described in the manufacturer’s protocol. Subsequently, libraries were size selected for 150-400 bp range with Pippen HT and size verified with Bioanalyzer. Samples were sequenced on Illumina NextSeq500 using 75bp paired-end reads to achieve sequencing depth of approximately 40M reads/sample.

### 10.2 CUT&RUN Bioinformatic Analysis

CUT&RUN paired-end data was trimmed using fastp v. 0.19.5 (Chen et al., 2018) to remove adapters and low quality base pairs (base pair quality score <30 and read length <35 bp). Trimmed reads were aligned to the hg38 human reference genome using bowtie2 v. 2.3.5 (Langmead and Salzberg, 2012) with the same alignment setting as described previously (Skene et al., 2018). Unaligned reads and discordantly aligned reads were eliminated and only primary aligned loci were kept. Duplicated reads were kept for the downstream analysis. The trimmed reads were also aligned to the S. cerevisiae yeast genome (sacCer3) yeast genome to evaluate the yeast spike-in reads for the spike-in normalization using the same bowtie2 parameters as described previously (Skene et al., 2018).

Control and miR-130a KD samples with triplicates were pooled and the peaks were called from the pooled bam files using MACS2 v. 2.2.5 (Zhang et al., 2008) with normalizing the samples using the yeast spike-in reads quantification. Calling peaks was performed by comparing the control and miR-130a KD CUT&RUN samples with the IgG and the peaks were filtered with the q-value cutoff <0.001. Peaks from different samples were intersected in a pairwise manner to determine the shared and unique peaks between two samples. Only shared peaks present with ETO and Flag antibodies in control and miR-130a KD samples were used for the downstream analysis. ChipPeakAnnot library (Zhu et al., 2010) in the R package v.3.6.1 was used to determine the overlap between peak sets and determine the shared and unique peaks represented in the Venn diagrams. Minimum peak overlap was set to 100 bp to determine the intersection peaks between the two samples. Peaks were annotated to determine the distribution across the genomic features such as promoters, 5’UTR, 3’UTR, introns, exons and intergenic regions using the Refseq genome annotation. Promoters were defined as the genomic regions starting 2Kb upstream of the transcriptional start site until 500 bp downstream of the transcriptional start site. Genomic distribution of the bound regions was represented in donut plots. P-values and odds ratios have been calculated for the counts of the peaks that were annotated for genomic regions for the miR-130a KD specific peaks compared with their counterparts in the shared peaks between control and miR-130a KD using Fisher’s exact test.

Motif analysis was performed on called peaks using homer v. 4.8 (Heinz et al., 2010). Normalized enrichment score (NES) for each motif was calculated as the fold change of the target to the background percentage. If the target percentage is less than 5, the pseudo count 1 was added to the target and the background percentages before calculating the fold change to attenuate the fold change of motifs with low target percentages. Volcano plots for the log10(p-value) vs the NES were plotted for each motif analysis. Only motives with NES>1.5 were displayed. Unique peaks to control and to miR-130a KD were clustered based on their signal intensity scores using K-means clustering algorithm. Peaks unique to Control were clustered into two clusters and peaks unique to miR-130a KD were clustered into three clusters. Motif analysis was performed on the clusters of the peaks using Homer.

Gene ontology analysis was performed on the list of genes that have AML1-ETO peaks bound to their promoters. PANTHER web server v. 15.0 (Thomas et al., 2003) was used for gene ontology analysis. Whole genome coverage tracks in bigwig file format were created from the bam files using the bamCoverage command in the deepTools package v.3.5.0 (Ramírez et al., 2016). The heatmaps of the signal in the Control and miR-130a KD samples for the peaks have been plotted from the coverage tracks using the computeMatrix and the plotHeatmap commands in the deepTools package.

Shared peaks which are common in ETO and Flag in the Control condition (Ctrl_ETO_Flag) have been compared with the previously published CUT&RUN data in Stengel et al. 2021. We have processed the Kasumi-1_(-)dTAG_anti-HA-1 and Kasumi-1_(-)dTAG_anti-HA-2 samples from the GSE153279 repository. The CUT&RUN samples have been downloaded, trimmed by fastp, and aligned to hg38 genome using bowtie2 aligner. Since there is no IgG sample provided in the repository for these samples, we called peaks using MACS2 without using control/background. We determined the intersection between our Ctrl_ETO_Flag peaks and the peaks of these two public data samples by allowing a minimum overlap of 100bp to determine the intersection between the peak sets.

### 10.3 CUT&RUN and RNA-seq Enrichment map

Thirty-one genes that have AML1-ETO peaks bound to their promoters and that are upregulated following miR-130a KD in Kasumi cells at FDR <0.05 have been subjected to pathway enrichment analysis testing against the GO biological process and MSigDB C2 databases using g:Profiler. Significant enrichment results under FDR 0.05 were visualized using Cytoscape/EnrichmentMap. For AML1-ETO regulated genes, publicly available datasets including AML1-ETO OE in human CD34^+^ HSPC (Tonks et al., 2007), AML1-ETO KD in Kasumi (Corsello et al., 2009) and dTAG AML1-ETO-FKBP12F36V degradation in Kasumi cells (Stengel et al., 2021). For AML1-ETO OE, genes upregulated and downregulated 3 days following AML1-ETO OE were used in the analysis (E-MEXP-583). For AML-ETO KD, differentially expressed genes following nucleofection of AML1-ETO-directed siRNAs were used in the analysis. For AML-ETO degradation, a high confidence AML1-ETO repression signature consisting of 59 genes was used in the analysis (<u>GSE153264</u>).

### Quantification and Statistical Analysis

The GraphPad Prism was used for all statistical analysis of xenografted mice and R was used for the bioinformatic analyses. Unless otherwise noted, mean ± SEM values are shown in the graphs. Statistical significance and p values were calculated with Mann-Whitney U-test or unpaired student’s t-test. Stars are used to indicate significance in figures: * p<0.05, ** p<0.01, *** p<0.001, *** p<0.0001.

### KEY RESOURCES TABLE (provided separately)

Supplemental Information includes Supplementary Figures 1-6 and Tables 1-5.

