## Supplemental Figures for "Identification of the Global miR-130a Targetome Reveals a Novel Role for TBL1XR1 in Hematopoietic Stem Cell Self-Renewal and t(8;21) AML"

**Supplemental Information**

- Supplementary Figures, Titles, Legends

**Figure S1.** miR-130a Overexpression and Knock-Down Alters Engraftment and Lineage Output of HSPC.
**Figure S2.** Enforced Expression of miR-130a Expands HSC by Forcing Them Into the Cell Cycle.
**Figure S3.** Mass Spectrometry and Chimeric AGO2 eCLIP Reveal miR-130a Targetome in Human HSPC.
**Figure S4.** Repression of TBL1XR1 Expands LT-HSC and Causes Differentiation Blocks.
**Figure S5.** Expression of miR-130a is Elevated in t(8;21) and its Loss of Function Cause Differentiation of Leukemia Cells
**Figure S6.** miR-130a KD Alters the Composition and Binding of AML1-ETO complex.

- Supplementary Tables

**Table S1**: Differentially Expressed Genes from RNA-seq of $CD34^+$ HSPC Following miR-130a OE, TBL1XR1 KD and Kasumi-1 cells Following miR-130a KD (Excel Table)
**Table S2**. Differentially Expressed Proteins from Mass Spectrometry Analysis of $CD34^+$ HSPC Following miR-130a OE (Excel Table)
**Table S3.** Quality Control Data for Chimeric AGO2 eCLIP
**Table S4.** miR-130a-Target Chimeras from AGO2 eCLIP in $CD34^+$ CB and Kasumi-1 cells (Excel Table)
**Table S5.** CBF AML Patient Information (Excel table)

#### Figure S1.

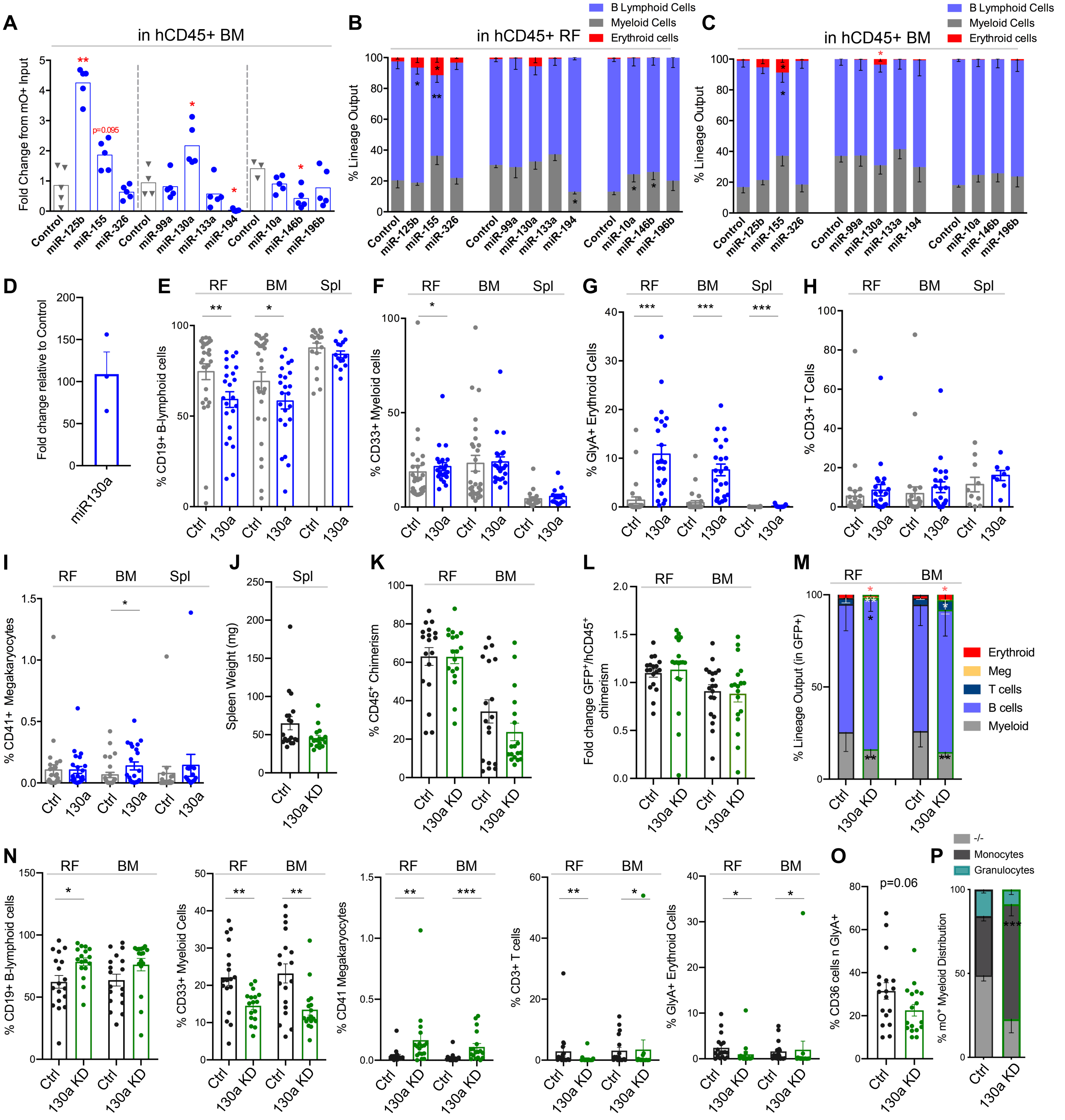

A
in hCD45+ BM
Fold Change from mO+ Input
p=0.095
Control
miR-125b
miR-155
miR-326
Control
miR-99a
miR-130a
miR-133a
miR-194
Control
miR-10a
miR-146b
miR-196b
B
in hCD45+ RF
B Lymphoid Cells
Myeloid Cells
Erythroid cells
% Lineage Output
C
in hCD45+ BM
B Lymphoid Cells
Myeloid Cells
Erythroid cells
% Lineage Output
D
Fold change relative to Control
miR130a
E
RF
BM
Spl
% CD19+ B-lymphoid cells
Ctrl
130a
F
% CD33+ Myeloid cells
G
% GlyA+ Erythroid Cells
H
% CD3+ T Cells
I
% CD41+ Megakaryocytes
J
Spl
Spleen Weight (mg)
Ctrl
130a KD
K
% CD45+ Chimerism
L
Fold change GFP+/hCD45+ chimerism
M
% Lineage Output (in GFP+)
Erythroid
Meg
T cells
B cells
Myeloid
N
% CD19+ B-lymphoid cells
% CD33+ Myeloid Cells
% CD41 Megakaryocytes
% CD3+ T cells
% GlyA+ Erythroid Cells
O
p=0.06
% CD36 cells n GlyA+
P
-/-
Monocytes
Granulocytes
% mO+ Myeloid Distribution

**Figure S1. miR-130a Overexpression and Knock-Down Alters Engraftment and Lineage Output of HSPC.**

(A) Fold change of $mO^+CD45^+$ cells in right femur (RF) at 24 weeks post-transplantation compared to input levels following enforced expression of individual miRNAs in HPSC.

(B) Lineage output of $mO^+CD45^+$ cells from RF of xenografted mice at 24 weeks.

(C) Lineage output of $mO^+CD45^+$ cells from BM of xenografted mice at 24 weeks.

(D) qRT-PCR of miR-130a expression levels in $mO^+$ $CD45^+$ cells from xenografts at 12 weeks (n=3, each replicate represents pooled RF and BM from 4-5 mice.

(E) Percentage of $CD19^+$ B lymphoid cells in $mO^+CD45^+$ cells from RF, BM and Spl of control and miR-130a OE xenografts.

(F) Percentage of $CD33^+$ myeloid cells in $mO^+$ $CD45^+$ cells from RF, BM and Spl.

(G) Percentage of $GlyA^+$ erythroid cells in $mO^+$ $CD45^-$ cells from RF, BM and Spl.

(H) Percentage of $CD3^+$ T lymphoid cells in $mO^+CD45^+$ cells from RF, BM and Spl.

(I) Percentage of $CD41^+$ megakaryocytes in $mO^+CD45^-$ cells from RF, BM and Spl.

(J) Spleen weight of control and miR-130a KD xenotransplanted mice at 24 weeks.

(K) Human $CD45^+$ chimerism in RF and BM at 24 weeks post-transplantation with HSPC transduced with miR-130a KD or control lentiviruses (n=2 biological experiments, 7-10 mice/experimental group).

(L) Fold change of $GFP^+CD45^+$ cells at 24 weeks post-transplantation compared to input levels.

(M) Lineage distribution of $GFP^+$ xenografts.

(N) Proportion of $CD19^+$ B cells, $CD33^+$ myeloid cells, $CD41^+$ megakaryocytes, $CD3^+$ T cells and $GlyA^+$ erythroid cells in $GFP^+$ xenografts at 24 weeks.

(O) Proportion of $CD36^+$ erythroid precursors in $GFP^+GlyA^+$ cells.

(P) Proportion of granulocytes and monocytes in $GFP^+CD33^+$ cells.

(A-P) Mann-Whitney test, all error bars indicate ± SEM, $^*p<0.05$, $^{**}p<0.01$, $^{***}p<0.001$, $^{****}p<0.0001$

**Figure S2.**

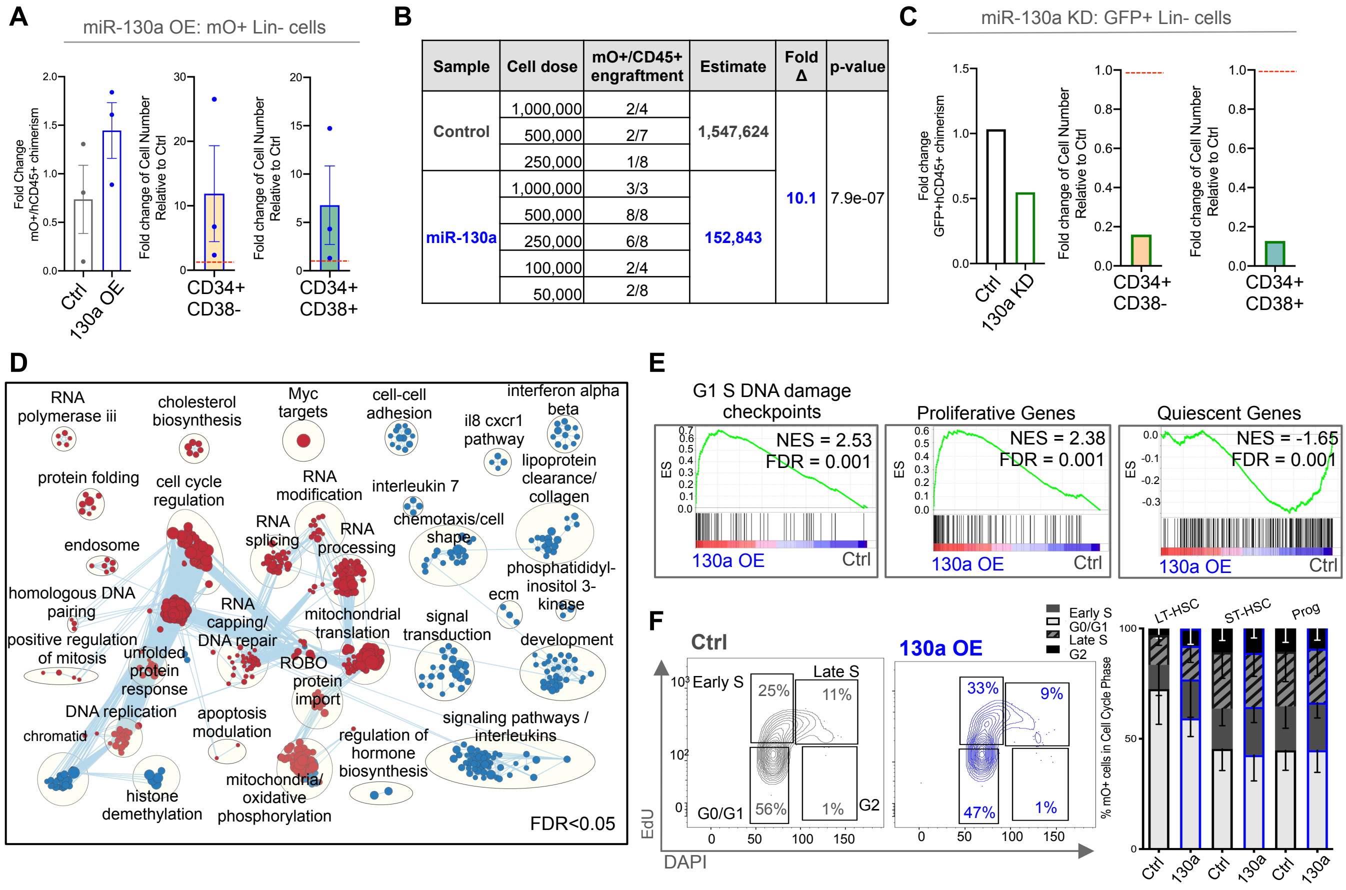


| Sample | Cell dose | mO+/CD45+ engraftment | Estimate | Fold Δ | p-value |
|---|---|---|---|---|---|
| Control | 1,000,000 | 2/4 | 1,547,624 | 10.1 | 7.9e-07 |
| | 500,000 | 2/7 | | | |
| | 250,000 | 1/8 | | | |
| miR-130a | 1,000,000 | 3/3 | 152,843 | | |
| | 500,000 | 8/8 | | | |
| | 250,000 | 6/8 | | | |
| | 100,000 | 2/4 | | | |
| | 50,000 | 2/8 | | | |

**Figure S2. Enforced Expression of miR-130a expands HSC by Forcing Them Into the Cell Cycle.**

(A) Fold change in the proportion of $mO^+CD45^+$ HSPC compared to input conditions (left), fold change of $mO^+CD34^+CD38^-$ (middle) and $mO^+CD34^+CD38^+$ (right) cell number in miR-130a OE mice relative to control.

(B) Table outlining cell doses and number of mice engrafted at each dose in the secondary NSG mice transplanted with $CD45^+mO^+$ cells from control and miR-130a OE xenografts.

(C) Fold change in the proportion of $GFP^+CD45^+$ HSPC compared to input conditions (left), fold change of $GFP^+CD34^+CD38^-$ (middle) and $GFP^+CD34^+CD38^+$ (right) cell number in miR-130a KD mice relative to control.

(D) Enrichment map of upregulated and downregulated gene sets in $mO^+CD34^+$HSPC following miR-130a OE. FDR<0.05, Mann Whitney p<0.05, node size is proportional to NES.

(E) GSEA plot showing enrichment of G1/S DNA damage checkpoints, proliferative and quiescence genes following miR-130a OE compared to control.

(F) Cell cycle and proliferation analysis of sorted LT-HSC, ST-HSC and progenitor cells 3 days post-transduction with control and miR-130a OE lentiviruses.

### Figure S3.

A

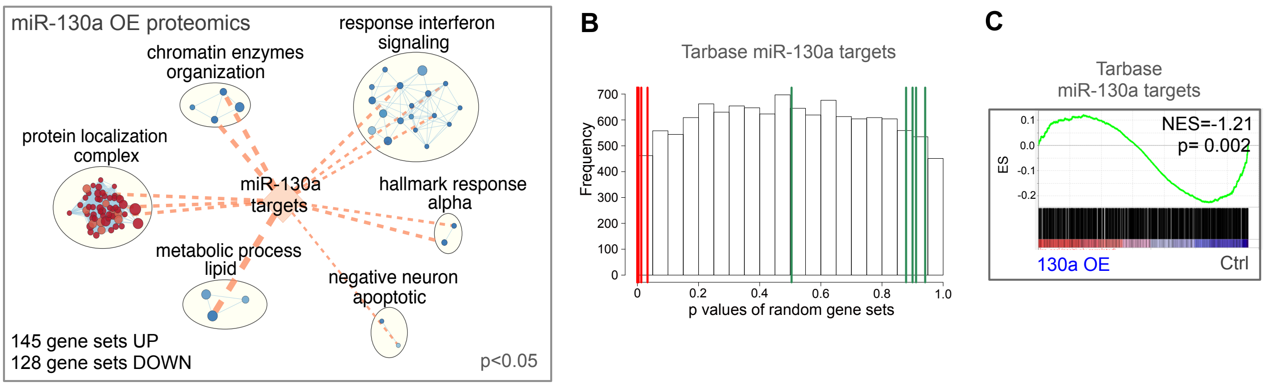


D

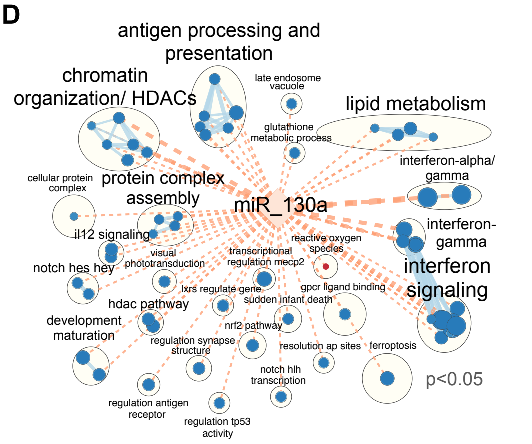


E

Targets downregulated by miR-130a OE

| gene | description | logFC | AveExpr | t | P.Value | adj.P.Val | miR-130a occurence |
|---|---|---|---|---|---|---|---|
| MTMR9 | myotubularin related protein 9 | -0.42 | 23.32 | -4.21 | 0.013 | 0.999 | 2 |
| TSC22D2 | TSC22 domain family member 2 | -0.93 | 23.37 | -4.03 | 0.015 | 0.999 | 1 |
| SLC35E1 | solute carrier family 35 member E1 | -1.30 | 21.60 | -3.95 | 0.027 | 0.999 | 1 |
| SAR1B | secretion associated Ras related GTPase 1B | -0.44 | 24.34 | -3.90 | 0.017 | 0.999 | 1 |
| JARID2 | jumonji and AT-rich interaction domain containing 2 | -0.46 | 22.73 | -3.59 | 0.022 | 0.999 | 4 |
| ANKRD52 | ankyrin repeat domain 52 | -0.46 | 24.61 | -3.56 | 0.022 | 0.999 | 7 |
| TBL1XR1 | transducin beta like 1 X-linked receptor 1 | -0.40 | 27.28 | -3.22 | 0.031 | 0.999 | 4 |
| TXNIP | thioredoxin interacting protein | -0.36 | 25.83 | -2.96 | 0.040 | 0.999 | 3 |
| DICER1 | dicer 1, ribonuclease III | -0.31 | 26.12 | -2.87 | 0.044 | 0.999 | 5 |
| CBFB | core-binding factor beta subunit | -0.31 | 27.03 | -2.75 | 0.050 | 0.999 | 2 |
| PSAP | prosaposin | -0.25 | 30.14 | -2.74 | 0.051 | 0.999 | 9 |
| TNRC6B | trinucleotide repeat containing 6B | -0.25 | 24.53 | -2.73 | 0.051 | 0.999 | 2 |
| VAMP3 | vesicle associated membrane protein 3 | -0.73 | 22.84 | -2.54 | 0.062 | 0.999 | 1 |
| PDZD11 | PDZ domain containing 11 | -0.36 | 22.61 | -2.46 | 0.068 | 0.999 | 3 |
| SECISBP2L | SECIS binding protein 2 like | -0.31 | 22.91 | -2.46 | 0.068 | 0.999 | 2 |
| NDUFA7 | NADH:ubiquinone oxidoreductase subunit A7 | -0.48 | 25.94 | -2.31 | 0.080 | 0.999 | 5 |
| VGLL4 | vestigial like family member 4 | -1.88 | 24.81 | -2.25 | 0.086 | 0.999 | 8 |
| RPA2 | replication protein A2 | -0.22 | 28.80 | -2.24 | 0.086 | 0.999 | 2 |
| ZBTB7A | zinc finger and BTB domain containing 7A | -0.26 | 25.16 | -2.24 | 0.087 | 0.999 | 1 |
| ATF7 | activating transcription factor 7 | -0.84 | 23.48 | -2.07 | 0.105 | 0.999 | 1 |

F

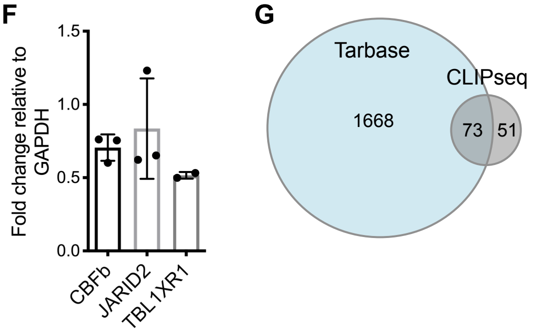


H

miR-130a eCLIPseq chimera targets in 3'UTR

Leading edge genes

| SYMBOL | RANK METRIC SCORE |
|---|---|
| KIT | -1.773 |
| TFB1M | -2.027 |
| ZNF3 | -2.046 |
| EFR3A | -2.257 |
| CTSA | -2.546 |
| TNRC6B | -2.726 |
| TBL1XR1 | -3.222 |
| JARID2 | -3.587 |
| MTMR9 | -4.214 |

I

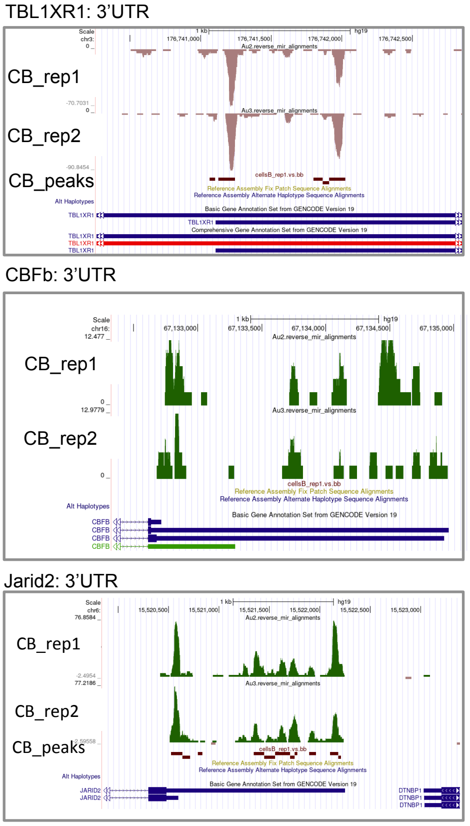


J

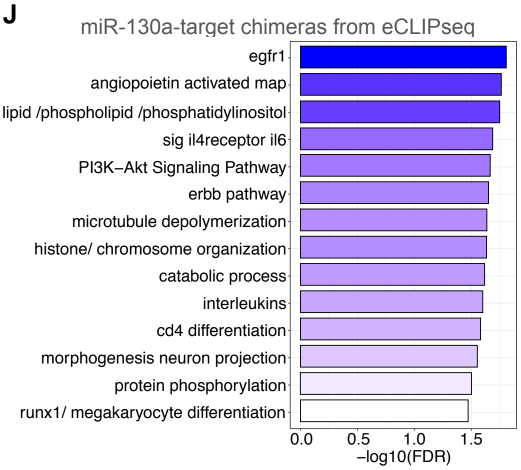

**Figure S3. Mass Spectrometry and Chimeric AGO2 eCLIP Reveal miR-130a Targetome in Human HSPC.**

(A) Enrichment map of gene sets containing upregulated (red) and downregulated (blue) proteins following miR-130a OE in $CD34^+$ CB cells, Wilcoxon two-sided test, $p<0.05$.

(B) Enrichment of Tarbase-predicted miR-130a targets in proteins downregulated in $CD34^+$ HSPC following miR-130a OE. Histogram represents the 1230 gene sets from miR-130a OE VS control and 2000 randomly selected gene sets from the HUGO gene list. Red lines represent p values for targets of 7 miRNAs from miR-130 family, green lines represent p values for targets of 7 randomly selected miRNAs.

(C) GSEA of Tarbase miR-130a targets showing a depletion of predicted targets in proteome changes following miR-130a OE.

(D) Enrichment map of miR-130a tarbase predicted targets in downregulated (blue) proteins in $CD34^+$ HSPC following miR-130a OE compared to control. Node size is proportional to NES; miR-130a tarbase targets used as a signature gene set; Wilcoxon one-sided test, $p<0.05$

(E) Table showing top 20 down regulated proteins in miR-130a OE cells compared to control that are also Tarbase predicted targets. The occurrence column states how many cell lines the gene was found as a target.

(F) Quantitative changes in protein levels of miR-130a targets detected by capillary-based western blot.

(G) Overlap of miR-130a targets from Tarbase and chimeric AGO2 CLIP-seq in $CD34^+$ CB cells.

(H) Leading edge analysis of miR-130a-target chimeras within proteins down regulated after miR-130a OE.

(I) UCSC genome browser tracks of chimeric AGO2 CLIP-seq reads showing peaks corresponding to miR-130a binding in the 3'UTR of TBL1XR1, CBFß and JARID2.

(J) Bar graph representing enriched gene sets in miR-130a-target chimeras from eCLIP.

**Figure S4.**

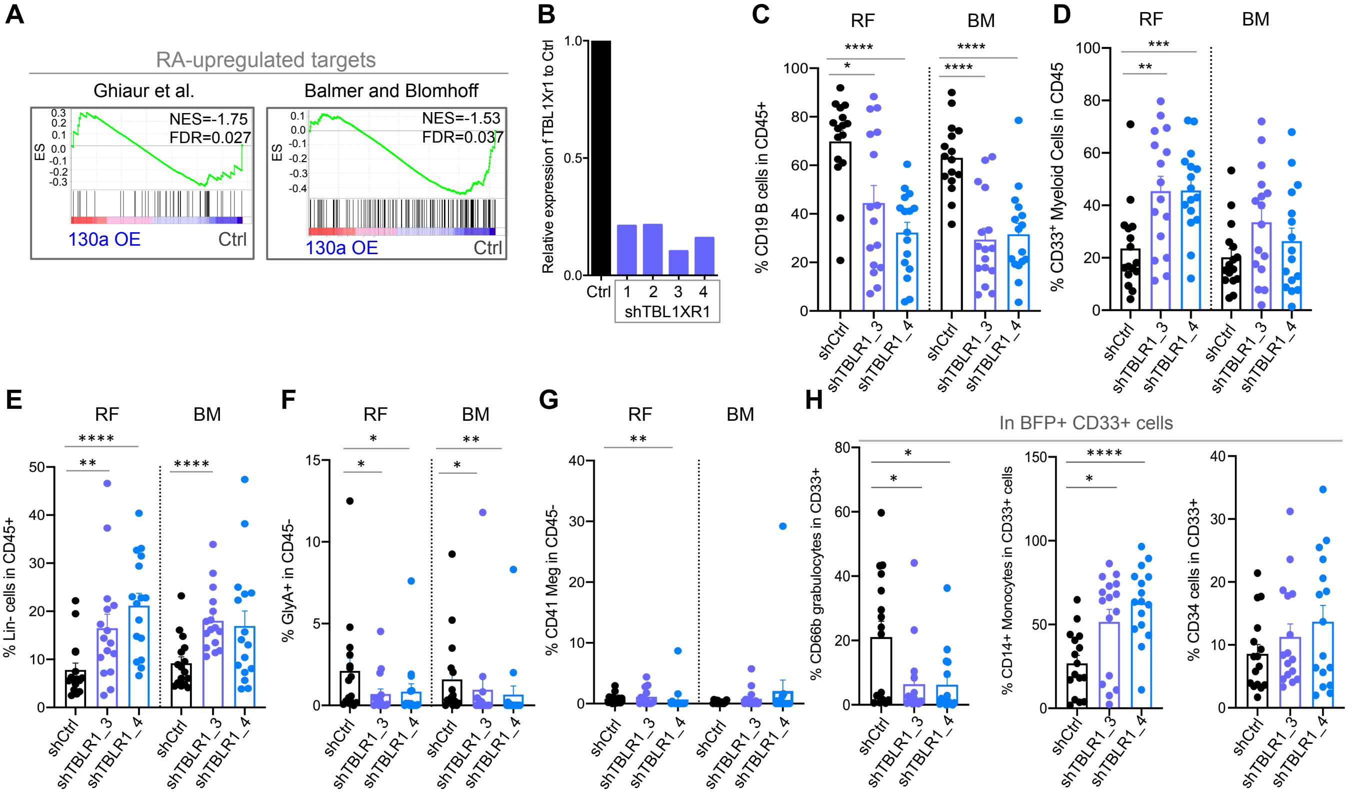


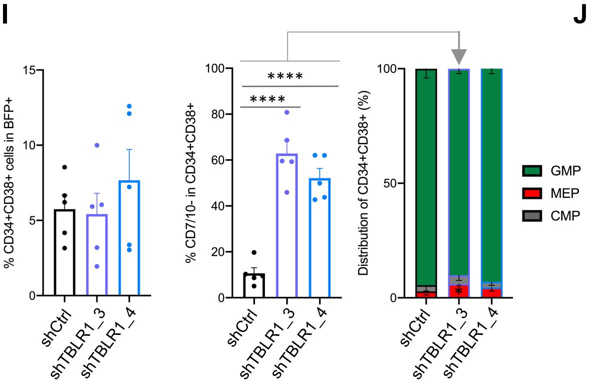


J

| Condition | Cell Dose | Engrafted Mice | Frequency | p Value | FC to Ctrl |
|---|---|---|---|---|---|
| shCtrl | 300K | 3/3 | 1:76,070 | | |
| | 100K | 3/4 | | | |
| | 30K | 1/4 | | | |
| TBL1XR1 sh3 | 300K | 1/1 | 1:16,229 | 0.01 | **4.7** |
| | 100K | 4/4 | | | |
| | 30K | 3/4 | | | |
| | 10K | 2/3 | | | |
| | 5K | 1/4 | | | |
| TBL1XR1 sh4 | 100K | 1/1 | 1:29,007 | 0.16 | **2.6** |
| | 30K | 1/3 | | | |
| | 10K | 2/4 | | | |
| | 5K | 1/4 | | | |

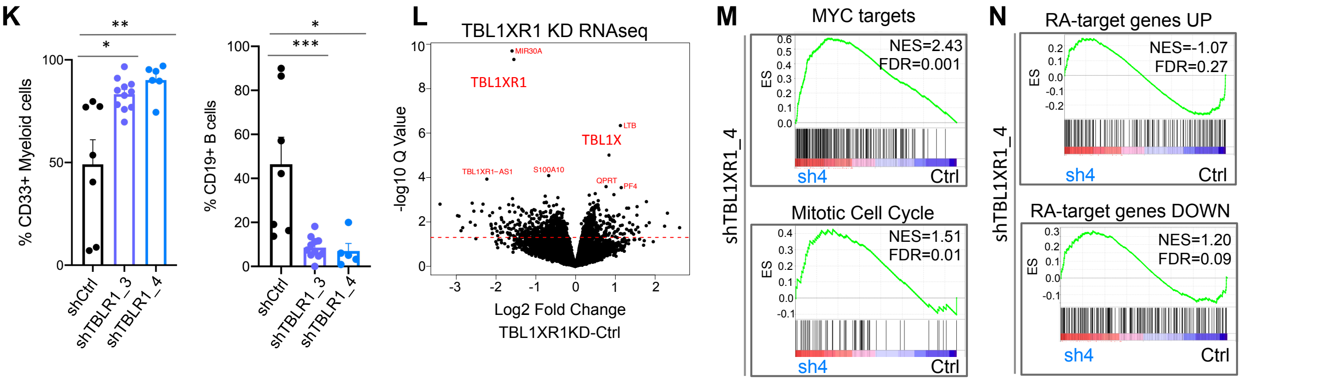

**Figure S4. Repression of TBL1XR1 Expands HSC and Causes Differentiation Blocks.**
(A) GSEA plots showing depletion of genes upregulated by RA in transcriptome profile following miR-130a OE in $CD34^+$ HSPC.
(B) TBL1XR1 mRNA levels measured by qRT-PCR in 293T cells 2 days post-transfection with 4 different shRNAs.
(C) Percentage of $CD19^+$ B lymphoid cells in $BFP^+CD45^+$ cells from control and TBL1XR1 KD xenografts at 24 weeks.
(D) Percentage of $CD33^+$ myeloid cells in $BFP^+CD45^+$ cells from RF and BM.
(E) Percentage of Lin- cells in $BFP^+CD45^+$ cells from RF and BM.
(F) Percentage of $GlyA^+$ erythroid cells in $BFP^+CD45^-$ cells from RF and BM.
(G) Percentage of $CD41^+$ megakaryocytes in $BFP^+CD45^-$ cells from RF and BM.
(H) Percentage of $CD66b^+$ granulocytes, $CD14^+$ monocytes and $CD34^+$ cells in $CD33^+CD45^+BFP^+$ cells from RF.
(I) Proportion of $BFP^+CD34^+CD38^-$ cells in RF and frequency of CMP, GMP and MEP cell populations in 24 week xenografts from two independent biological experiments (n=5, each replicate contains pooled RF from 2-4 individual mice), unpaired t-test.
(J) Table outlining cell doses and number of mice engrafted at each dose in the secondary NSG-GF mice transplanted with $CD45^+BFP^+$ cells from control and TBL1XR1 KD xenografts.
(K) Percentage of $CD33^+$ myeloid and $CD19^+$ B lymphoid cells in RF and BM of secondary NSG-GF mice transplanted with control and TBL1XR1 KD cells.
(L) Volcano plot showing differentially expressed transcripts following TBL1XR1 KD in $CD34^+CD38^-$ CB cells.
(M) GSEA plots showing enrichment of MYC targets and mitotic cell cycle genes in transcriptome profile of $CD34^+CD38^-$ CB cells following TBL1XR1 KD.
(N) GSEA plots showing enrichment of RA-target genes in transcriptome profile of $CD34^+CD38^-$ CB cells following TBL1XR1 KD.
(C-H, K) Mann-Whitney test, all error bars indicate ± SEM, *$p<0.05$, **$p<0.01$, ***$p<0.001$, ****$p<0.0001$

### Figure S5

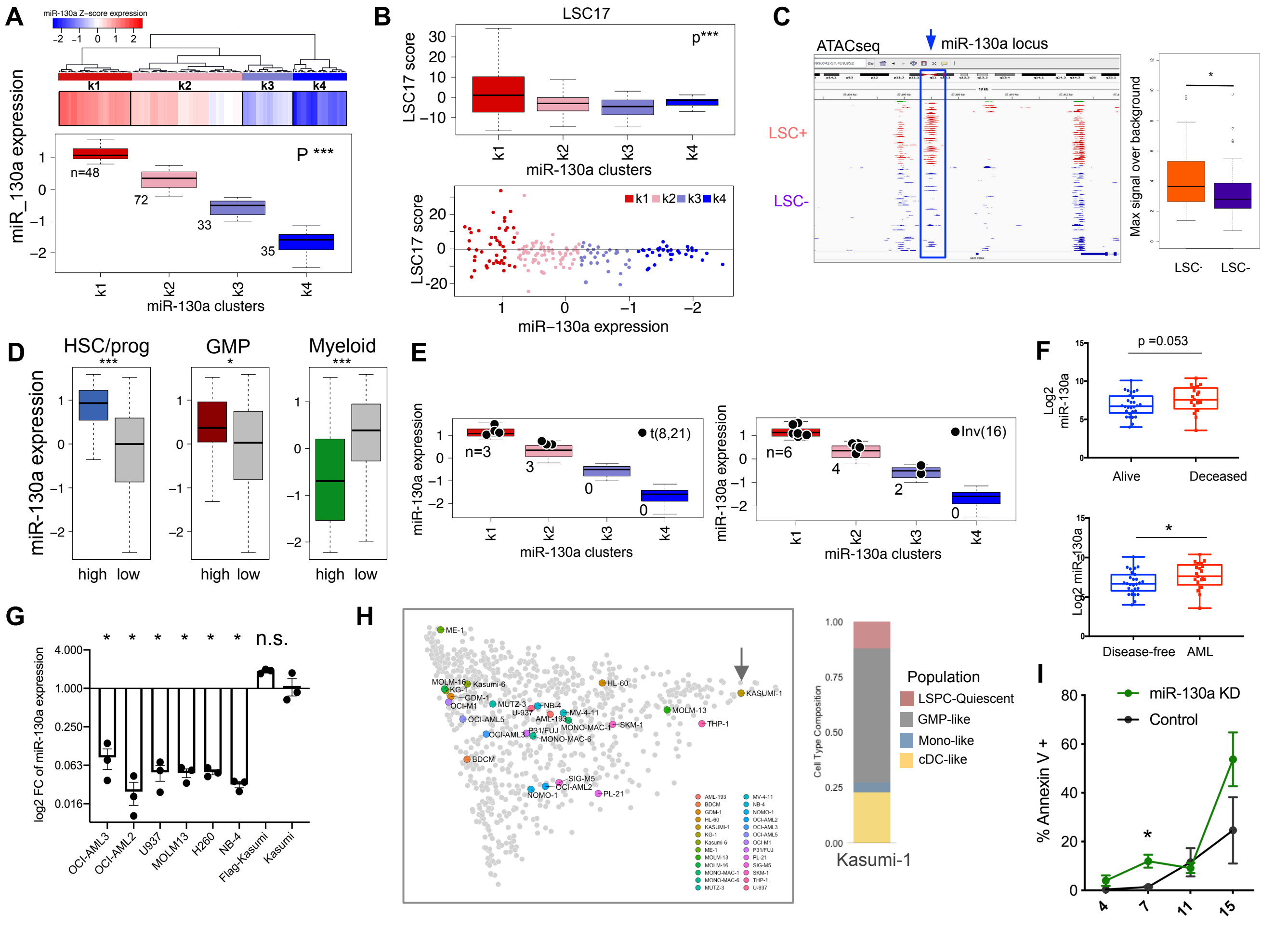


## J

| AML | Sample ID | Cytogenetics | Age | Gender | miR-130a FC |
|---|---|---|---|---|---|
| I | 100391 | 45,X,-Y,t(8;21)(q22;q22)[17]/ 46,XY[3] & 8;21 | 21 | male | 1.7 |
| II | 130307 | 8;21 & 46,XY,t(8;21) (q22;q22)[3]/46,XY[1] | 85 | male | 2.5 |
| III | 110841 | 45,X,-Y,t(8;21)(q22;q22)[10] & 8;21 | 29 | male | 1.6 |
| IV | 7625 | 46,XY,t(8;21)(q22;q22),del(9) (q13q22)[9]/46,XY[1] | 35 | male | 2.6 |
| V | 160036 | 8;21 & 45,X,-X,t(8;21) (q22;q22)[10] | 18 | female | 1.5 |
| IV | 090765 | 46,XX,t(8;21)(q22;q22)[8] | 95 | female | 1.3 |

#### L miR-130a KD and targets from eCLIP in Kasumi-1

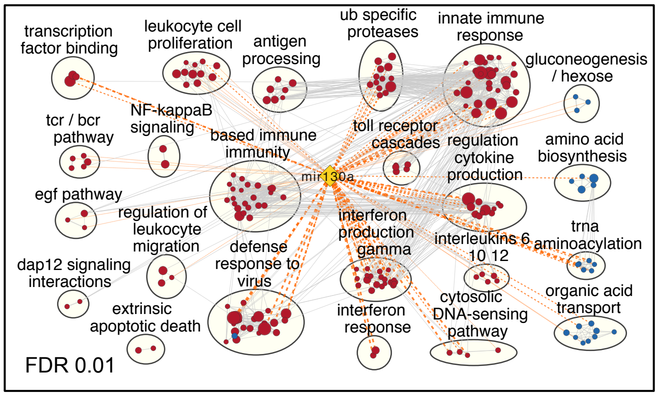


## K

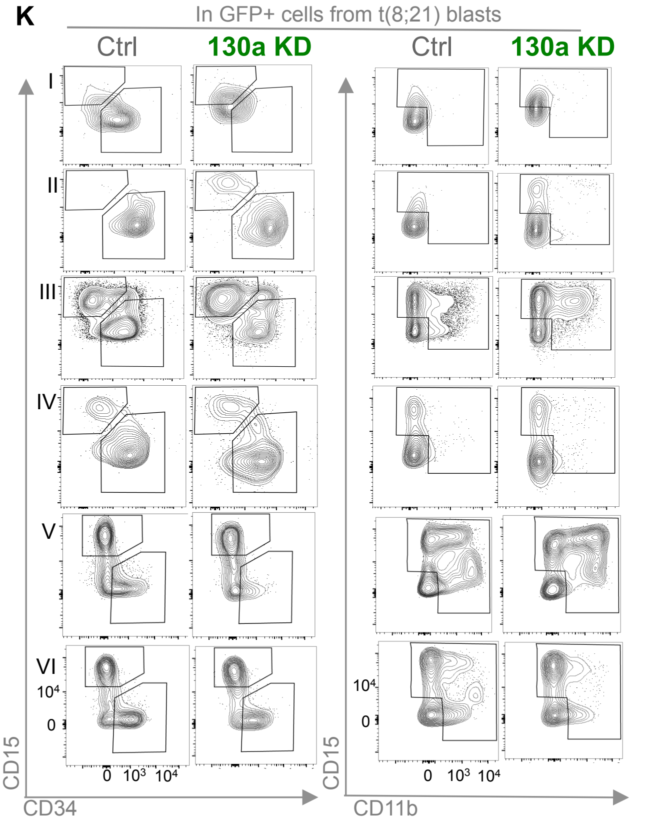

**Figure S5. Expression of miR-130a is Elevated in t(8;21) and its Loss of Function Causes Differentiation of Leukemia Cells**

(A) Clustering of TCGA AML based on miR-130a expression, ANOVA test, ***$p<0.001$.

(B) LSC17 score in the clusters ranked according to miR-130a levels.

(C) Chromatin accessibility surrounding miR-130a locus in sorted and functionally defined $LSC^+$ and $LSC^-$ AML fractions from primary AML samples.

(D) GSVA showing association of miR-130a levels with HSC/progenitor, GMP-like and Myeloid-like signatures using scRNAseq data from TCGA AMLs, Student's t-test, *$p<0.05$, ***$p<0.001$.

(E) Distribution of t(8;21) and inv(16) AMLs in the clusters based on miR-130a levels.

(F) Expression of miR-130a measured by microarray in CBF AML patient cohort, one tailed t-test, *$p<0.05$.

(G) qRT-PCR for miR-130a expression level in AML cell lines. RNU48 was used as an endogenous control, unpaired t-test, all error bars indicate ± SEM, *$p<0.05$.

(H) tSNE plots showing AML clustering and position of different cell lines among the clusters (left) and deconvolution of cell type composition of Kasumi-1 cells (right).

(I) Proportion of $annexinV^+$ apoptotic cells in Kasumi-1 cells transduced with control or miR-130a KD lentiviruses, unpaired t-test, *$p<0.05$.

(J) Table outlining t(8;21) AML patient information used in the *in vitro* assays.

(K) Flow cytometry plots presenting proportion of $CD34^+$, $CD15^+$ and $CD11b^+$ cells in $GFP^+$ t(8;21) AML blasts transduced with control of miR-130a KD lentiviruses.

(L) Enrichment map showing upregulated and downregulated gene sets following miR-130a KD in Kasumi and miR-130a targets from chimeric AGO2 cCLIP-seq, FDR<0.01, Mann Whitney, $p<0.05$.

Figure S6

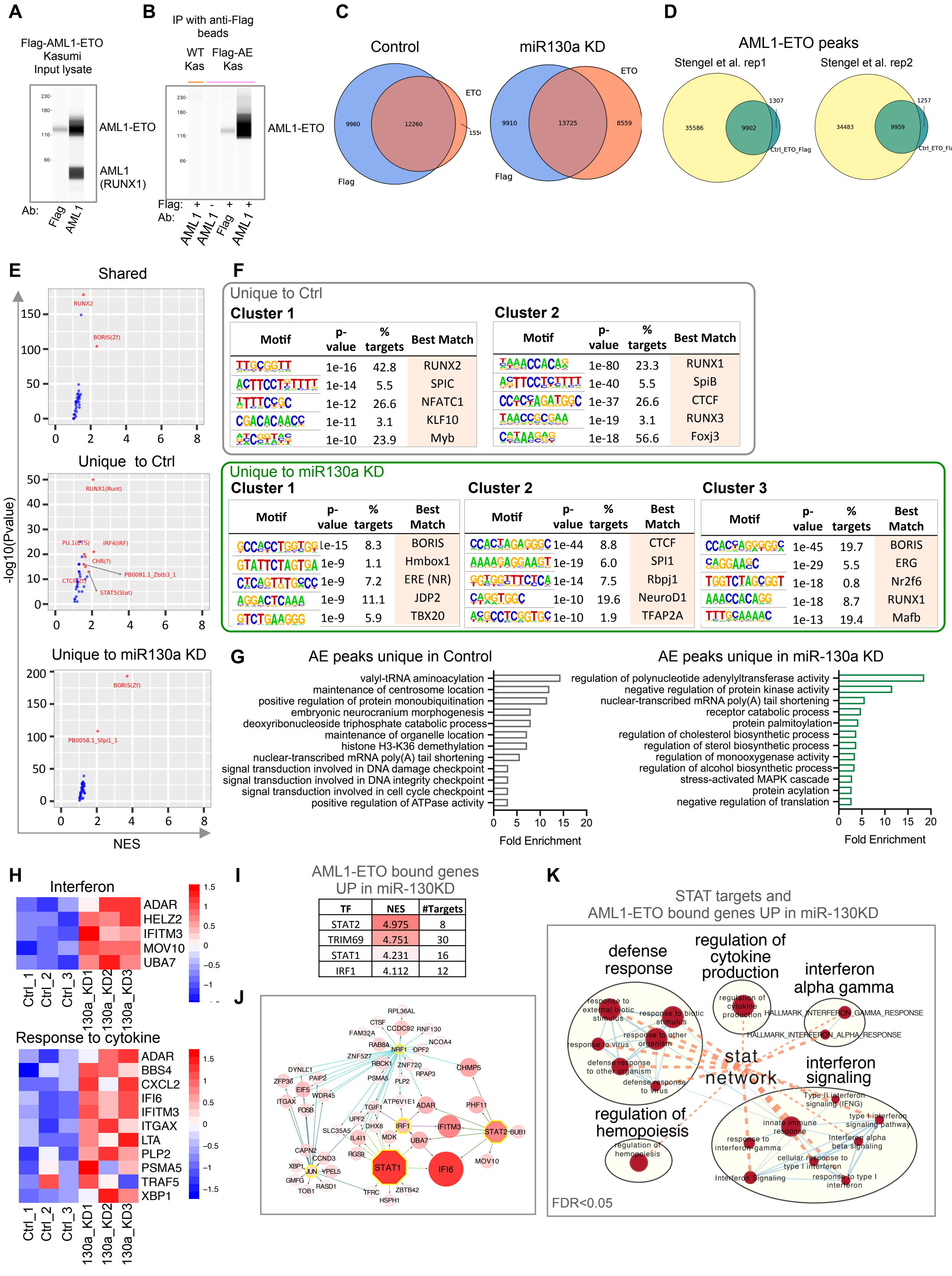

A
Flag-AML1-ETO Kasumi Input lysate
AML1-ETO
AML1 (RUNX1)
Ab: Flag AML1
B
IP with anti-Flag beads
WT Kas
Flag-AE Kas
AML1-ETO
Flag: + - + +
Ab: AML1 AML1 Flag AML1
C
Control
miR130a KD
ETO
Flag
9960
12260
9910
13725
8559
D
AML1-ETO peaks
Stengel et al. rep1
Stengel et al. rep2
35586
9902
1307
Ctrl_ETO_Flag
34483
9959
1257
E
Shared
RUNX2
BORIS(Zf)
Unique to Ctrl
RUNX1(Runt)
PU.1(ETS)
IRF4(IRF)
CHR(?)
PB0091.1_Zbtb3_1
CTCF(Zf)
STAT5(Stat)
Unique to miR130a KD
BORIS(Zf)
PB0058.1_Sfpi1_1
-log10(Pvalue)
NES
F
Unique to Ctrl
Cluster 1
Motif | p-value | % targets | Best Match
1e-16 | 42.8 | RUNX2
1e-14 | 5.5 | SPIC
1e-12 | 26.6 | NFATC1
1e-11 | 3.1 | KLF10
1e-10 | 23.9 | Myb
Cluster 2
Motif | p-value | % targets | Best Match
1e-80 | 23.3 | RUNX1
1e-40 | 5.5 | SpiB
1e-37 | 26.6 | CTCF
1e-19 | 3.1 | RUNX3
1e-18 | 56.6 | Foxj3
Unique to miR130a KD
Cluster 1
Motif | p-value | % targets | Best Match
1e-15 | 8.3 | BORIS
1e-9 | 1.1 | Hmbox1
1e-9 | 7.2 | ERE (NR)
1e-9 | 11.1 | JDP2
1e-9 | 5.9 | TBX20
Cluster 2
Motif | p-value | % targets | Best Match
1e-44 | 8.8 | CTCF
1e-19 | 6.0 | SPI1
1e-14 | 7.5 | Rbpj1
1e-10 | 19.6 | NeuroD1
1e-10 | 1.9 | TFAP2A
Cluster 3
Motif | p-value | % targets | Best Match
1e-45 | 19.7 | BORIS
1e-29 | 5.5 | ERG
1e-18 | 0.8 | Nr2f6
1e-18 | 8.7 | RUNX1
1e-13 | 19.4 | Mafb
G
AE peaks unique in Control
valyl-tRNA aminoacylation
maintenance of centrosome location
positive regulation of protein monoubiquitination
embryonic neurocranium morphogenesis
deoxyribonucleoside triphosphate catabolic process
maintenance of organelle location
histone H3-K36 demethylation
nuclear-transcribed mRNA poly(A) tail shortening
signal transduction involved in DNA damage checkpoint
signal transduction involved in DNA integrity checkpoint
signal transduction involved in cell cycle checkpoint
positive regulation of ATPase activity
Fold Enrichment
AE peaks unique in miR-130a KD
regulation of polynucleotide adenylyltransferase activity
negative regulation of protein kinase activity
nuclear-transcribed mRNA poly(A) tail shortening
receptor catabolic process
protein palmitoylation
regulation of cholesterol biosynthetic process
regulation of sterol biosynthetic process
regulation of monooxygenase activity
regulation of alcohol biosynthetic process
stress-activated MAPK cascade
protein acylation
negative regulation of translation
Fold Enrichment
H
Interferon
ADAR
HELZ2
IFITM3
MOV10
UBA7
Ctrl_1 Ctrl_2 Ctrl_3 130a_KD1 130a_KD2 130a_KD3
Response to cytokine
ADAR
BBS4
CXCL2
IFI6
IFITM3
ITGAX
LTA
PLP2
PSMA5
TRAF5
XBP1
I
AML1-ETO bound genes UP in miR-130KD
TF | NES | #Targets
STAT2 | 4.975 | 8
TRIM69 | 4.751 | 30
STAT1 | 4.231 | 16
IRF1 | 4.112 | 12
J
NRF1
IRF1
STAT2
STAT1
JUN
IFI6
IFITM3
ADAR
UBA7
K
STAT targets and AML1-ETO bound genes UP in miR-130KD
defense response
regulation of cytokine production
interferon alpha gamma
stat network
interferon signaling
regulation of hemopoiesis
HALLMARK_INTERFERON_GAMMA_RESPONSE
HALLMARK_INTERFERON_ALPHA_RESPONSE
FDR<0.05

**Figure S6. miR-130a KD Alters the Composition and Binding of AML1-ETO complex.**

(A) Western blot of Flag-tagged AML-ETO in Kasumi-1 cell line using anti-Flag and anti-AML1 antibodies.

(B) Western blot of immunoprecipitated Flag-tagged AML1-ETO in wild-type and Flag-AML1-ETO knock-in Kasumi-1 cells with anti-Flag and anti-AML1 antibodies.

(C) Overlap of AML1-ETO peaks detected by anti-Flag and anti-ETO antibodies in Kasumi-1 cells transduced with control and miR-130a KD lentiviruses. Minimum distance of 100 bp was considered for peak overlap, q value $<0.001$.

(D) Overlap of AML1-ETO peaks published by Stengel et al. and peaks from our datasets.

(E) HOMER transcription factor binding site motif enrichment analysis of shared and unique peaks from control and miR-130a KD Kasumi-1 cells. Transcription factor motives with NES $>1.5$ are shown.

(F) K means clustering of AML1-ETO peaks unique to control and miR-130a KD.

(G) Gene Ontology Pathway analysis for shared and unique peaks in control and miR-130a KD Kasumi-1 cells using promoter-bound AML1-ETO genes.

(H) Heat maps showing genes in interferon and cytokine response pathways enriched upregulated, promoter-bound AML1-ETO genes unique to miR-130a KD.

(I) Iregulon transcription factor enrichment analysis of upregulated, promoter-bound AML1-ETO genes unique to miR-130a KD.

(J) STAT network showing enrichment of STAT target genes in upregulated, promoter-bound AML1-ETO genes unique to miR-130a KD.

(K) Enrichment map showing STAT targets among the upregulated, promoter-bound AML1-ETO genes unique to miR-130a KD, FDR$<0.05$, Mann-Whitney 2-sided test, $p<0.001$.

**Table S3. Quality Control Data for Chimeric AGO2 eCLIP**. PCR cycle number, library yield, number of raw, mapped, chimeric reads and significant peaks in global and miR-130a targeted Chimeric AGO2 eCLIP and size-matched inputs libraries from $CD34^{+}$ CB cells and Kasumi-1 cells.

| Sample | PCR cycle number | Library yield (fmoles) | Sample ID | Raw reads | Uniquely mapped reads (after removing repetitive reads) | PCR-deduplicated usable reads (total) | PCR-deduplicated usable reads (chimeric) | Significant peaks |
|---|---|---|---|---|---|---|---|---|
| Kasumi-1 total chimeric eCLIP | 14 | 123 | Au1 | 62,838,139 | 27,206,530 | 15,429,174 | 503,654 | 10,675 |
| Kasumi-1 total chimeric eCLIP size-matched input | 8 | 107 | Au5 | 49,869,528 | 13,581,963 | 11,640,962 | n/a | n/a |
| CD34+ CB miRNA total chimeric eCLIP, rep 1 | 14 | 85 | Au2 | 36,085,496 | 11,904,175 | 6,569,500 | 400,737 | 10,957 |
| CD34+ CB miRNA total chimeric eCLIP size-matched input, rep 1 | 8 | 167 | Au6 | 29,286,571 | 8,127,314 | 7,155,100 | n/a | n/a |
| CD34+ CB miRNA total chimeric eCLIP, rep 2 | 14 | 53.9 | Au3 | 32,429,965 | 11,145,131 | 6,805,423 | 385,270 | 11,009 |
| CD34+ CB miRNA total chimeric eCLIP size-matched input, rep 2 | 8 | 190 | Au7 | 26,374,635 | 7,793,585 | 7,097,389 | n/a | n/a |
| Kasumi-1 hsa-miR-130a targeted chimeric eCLIP | 13+12 | 239 | Au9 | 7,211,775 | 2,344,167 | n/a | 53,588 | n/a |
| CD34+ CB hsa-miR-130a targeted chimeric eCLIP | 13+12 | 302 | Au13 | 9,445,872 | 3,273,159 | n/a | 49,435 | n/a |
